# CARs are organized in nanodomains in the plasma membrane of T cells that accumulate at tumor contact sites

**DOI:** 10.1101/2023.07.19.549702

**Authors:** Christina Verbruggen, Leon Gehrke, Nicole Banholzer, Arindam Ghosh, Sebastian Reinhard, Justus Weber, Sören Doose, Hermann Einsele, Michael Hudecek, Thomas Nerreter, Markus Sauer

## Abstract

Chimeric antigen receptors (CARs) are synthetic immune receptors that are expressed in T cells through genetic engineering. CAR-T cells have been successfully used to eradicate very advanced leukemias and lymphomas and their functional properties have been intensively studied. However, relatively little is known about the spatiotemporal expression and organization of CARs on the T-cell membrane and how this influences their efficacy. Here, we applied super-resolution microscopy to visualize CD19-, ROR1-, and ROR2-specific CARs in human CD4^+^ and CD8^+^ T cells that were engineered with lentiviral and transposon-mediated gene transfer. Our data show that the majority of CARs is organized in nanodomains virtually independent of the T cell type, CAR construct and expression level. Quantitative analyses revealed a slightly higher CAR density in transposon-engineered T cells correlating with higher antigen sensitivity and faster resolution of anti-tumor functions compared to lentivirally-engineered T cells. Live-cell fluorescence imaging revealed that both, CAR nanodomains and CAR monomers accumulate at tumor contact sites and form multifocal immunological synapses. Our study provides novel insights into the membrane organization of CARs with single-molecule resolution and illustrates the potential of advanced microscopy to inform the rational design of synthetic immune receptors for applications in immune cell therapy.

## Introduction

CAR-T cells are engineered T cells redirected to target an extracellular surface-antigen on tumor cells that is ideally exclusively expressed on malignant cells and not on healthy tissue^1, 2^, since even very low amounts of antigens can trigger CAR-T cell mediated cytotoxicity^3^. CAR-T cell therapy has already brought about a revolution in the treatment of hematological malignancies^4^. With an increasing number of target antigens under investigation for CAR-based therapies, the potential application spectrum of CAR-T cell therapy is steadily expanding^5, 6^, and substantial efforts in the field aim to make CAR-T cell therapy conceivable as a prospective therapeutic option in solid tumors and autoimmune disorders as well^4, 7, 8^.

In order to further improve the efficacy of CAR-T cell immunotherapy, predict therapeutic outcome and ensure optimal care for patients, there is a desire in the field to elucidate mechanistic details of CAR-T cell mediated anti-tumor activity and enable evaluation of the potency and safety of CAR-T cell products already at early stages of product development. At present, *in vitro* cytotoxicity assays, immunophenotyping, proliferation and cytokine secretion assays, and re-stimulation assays, as well as elaborate *in* and *ex vivo* strategies such as tumor-on-a-chip are used to evaluate the efficacy and safety of CAR-T cells at the preclinical level^9, 10^. Even though CAR-T cells are considered to employ TCR signaling^11, 12^, knowledge about the expression and molecular organization of CARs on the surface of lymphocytes and during their interactions with tumor cells remains limited^13^. So far, flowcytometric methods have been used to determine the expression of CARs by staining T cells with specific ligands or antibodies conjugated with fluorophores^14^.

Most commonly, indirect labeling methods that rely on co-expression of the CAR and a transduction marker such as LNGFR, Her2t or EGFRt are applied as a surrogate marker for CAR expression^15^. Several alternatives have been used, but all have disadvantages as well: synthetic peptide-based tags have been incorporated in the CAR receptor itself, such as c-myc or FLAG-tags^16^ that can potentially promote immunogenicity, or reagents targeting the CAR binding domain, e.g. CD19/Fc fusion proteins^17^, anti-CAR-scFv-idiotype antibodies^18^, or protein L^14^. The main limitation of these labeling techniques is that they cannot be applied to a multitude of different CARs, depend on binder affinity and can potentially affect the CAR construct integrity or CAR-T cell function. Furthermore, flow cytometry exhibits only limited sensitivity and does not permit the visualization of the distribution of CARs on the cell surface^3^. So far CAR-T cell imaging has been performed by standard fluorescence imaging methods that do not allow for real-time three-dimensional (3D) visualization of the interaction of tumor and CAR-T cells and that do not provide the required sensitivity and spatial resolution to quantify the expression and resolve the molecular organization of CARs in the plasma membrane of T cells^19–21^.

In recent years, single-molecule localization microscopy methods such as photoactivated localization microscopy (PALM)^22^ and *direct* stochastic optical reconstruction microscopy (*d*STORM)^23^ have been used successful to map plasma membrane constituents or associated proteins in unprecedented detail. For example, *d*STORM quantified the expression level of CD19 and CD20 on primary tumor cells required for CAR-T cell therapy^3^ and enabled the development of T cell signaling models in which TCR-signaling complexes are assembled by rearrangement of nanoscale protein nanodomains within the plasma membrane^24–27^. Furthermore, diverse nanoscale organizations of TCR-based immunological synapses have been reported in previous studies, one such example being the formation of multifocal immunological synapses consisting of several TCR clusters^28^. However, corresponding functional implications of these observed variations in the molecular architecture of immunological synapses remain largely unknown^28–30^. In relevance to CAR-T cells, recent reports suggest that the molecular structure, organization, and quality of the immunological synapse can predict CAR-modified cell efficacy and toxicity, which in turn, has significant implications in clinical outcomes^31, 32^. This highlights the relevance of high-resolution CAR imaging combined with functional analysis to advance mechanistic knowledge and enable informative early product screening.

Here we present a way of direct CAR detection in a receptor-intrinsic manner without inclusion of additional tags. The microscopic approach used allowed us to visualize the CAR on the T cell surface at single-molecule resolution and enabled the precise quantification of CAR expression. We found that our staining method is reliable on Jurkat cells as well as primary CD4^+^ and CD8^+^ T cells and optimized it for CD19-, ROR1-, and ROR2-specific CAR constructs. *d*STORM imaging revealed that the majority of expressed CARs forms nanodomains in the plasma membrane independent of CAR construct, T cell type, transgene delivery method and expression level. Co-incubation of CD19-specific CAR-T cells expressing CAR-GFP with tumor cells displayed a strong accumulation of CAR monomers and nanodomains at contact sites to tumor cells, with multiple interaction foci indicating the formation of a multifocal immunological synapse. Complementary experiments with CAR-T cells expressing ZAP70-GFP and tumor cells or donor-matched primary B cells confirmed the formation of multifocal immunological synapses.

## Results

### Quantification of plasma membrane CARs by *d*STORM

We first focused our attention on the quantification of the expression and lateral organization of CARs in the plasma membrane of CAR-T cells generated by lentiviral transduction (LV) or nucleofection using the Sleeping Beauty (SB) transposon system. We then verified our results with CAR constructs of various designs and specificities, targeting established cancer-immunotherapy targets such as CD19 and ROR1, as well as the novel antigen ROR2.

For fluorescence imaging and quantification of CARs in the plasma membrane of T cells, we used single-molecule sensitive super-resolution imaging by *d*STORM^23, 33^. To extract quantitative information from single-molecule localization data, the average number of blink events (localizations) measured for a single fluorescently labeled probe (i.e. antibody) can be used to determine the number of probes bound to CARs^3, 34, 35^. We verified our approach by detecting CD19-specific CARs expressed on Jurkat cells by lentiviral transduction using an Alexa Fluor 647 (AF647)-labeled antibody recognizing the linker connecting scFv heavy and light chains. *d*STORM imaging of cells was performed by total internal reflection fluorescence microscopy to selectively image the basal plasma membrane at high signal-to-noise ratio (Extended Data Fig. 1a and Supplementary Fig. 1a-d). To ensure saturation of accessible antigen epitopes on the plasma membrane we titrated the antibody concentration in separate experiments and used saturated antibody concentrations of 5 µg ml^-1^ in all *d*STORM quantification experiments (Extended Data Fig. 1b,c and Supplementary Fig. 1e-h).

First, we investigated the expression and distribution of CD19- and ROR1-specific CARs in the plasma membrane of primary CD4^+^ and CD8^+^ cells introduced by SB transposition. On CD4^+^ and CD8^+^ CD19-specific CAR-T cells we detected (128 ± 68) and (126 ± 72) localizations µm^-2^ (mean ± standard deviation), respectively (Fig. 1a-c). Primary CD4^+^ and CD8^+^ T cells expressing ROR1-specific CARs showed slightly higher localization densities of (135 ± 74) and (181 ± 71) µm^-2^, respectively (Supplementary Fig. 2). These data demonstrate that the CAR density can vary slightly for different CAR constructs and T cell phenotypes. Furthermore, all *d*STORM images revealed the existence of smaller and larger localization clusters in the plasma membrane, presumably translating to single CAR molecules and oligomeric co-localizations of several CAR molecules, which we term CAR monomers and nanodomains, respectively (Fig. 1, Extended Data Fig. 1, Supplementary Fig. 2 and Supplementary Fig. 3).

**Fig. 1.**
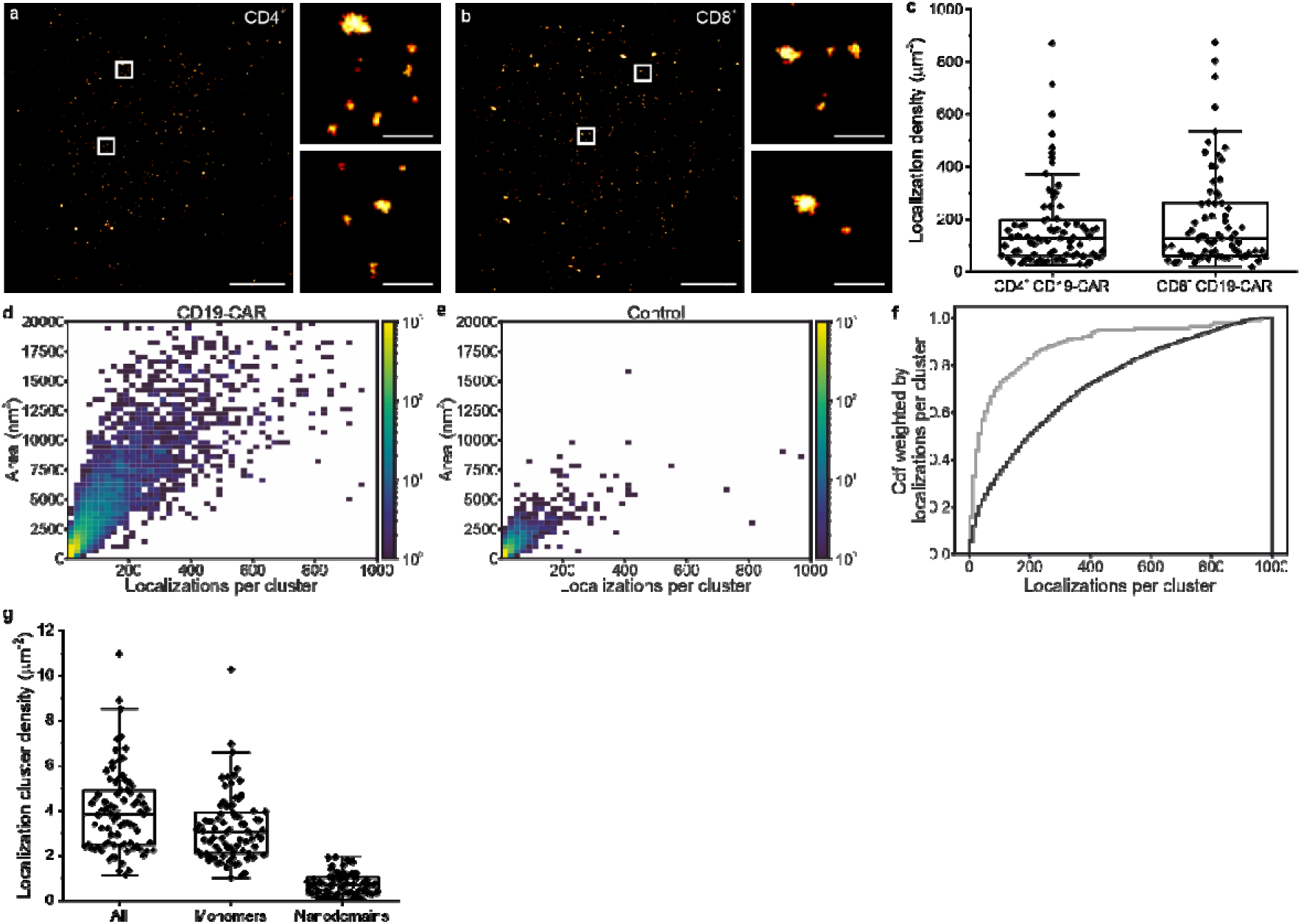
*d*STORM enables quantification of CAR expression and organization as monomers and in nanodomains in the plasma membrane of native CD4^+^ and CD8^+^ T cells. **a,b,** *d*STORM images of the basal plasma membrane of CD19-specific CARs on CD4^+^ (a) and CD8^+^ T cells (b). Small panels display magnifications of boxed regions revealing the partial organization of CARs in nanodomains in the plasma membrane independent of the T cell phenotype. Scale bars, 3 μm, 300 nm (magnified images). **c**, Boxplots show localization densities (localizations µm^-2^) of 128 ± 68 (CD4^+^), and 126 ± 72 (CD8^+^) for CD19-specific CARs (n=74-82 cells). **d,e**, Bivariate histogram from spatially unresolvable localization clusters in *d*STORM experiments using AF647 labeled antibodies staining primary T cells that do (CD19-CAR, d) or do not (control, e) express CD19-specific CARs. Histograms display the distribution of localizations per cluster and convex hull areas of localization clusters. **f**, Cumulative histogram of localizations per cluster weighted by the localizations per cluster for CD19-CAR (black) and control (gray). The plot reflects the percentage of CD19-specific CAR labels found within clusters with equal or less than the indicated localizations per cluster. All histograms were assembled from n=127 cells and N∼17,100 clusters for CD19-specific CARs and from n=124 cells and N∼4,800 clusters for the control. **g**, Localization cluster densities determined from the histogram shown in (d) for all localization clusters, small localization clusters (predominantly monomeric CARs with <40 localizations) and nanodomains (>40 localizations).

#### CARs form nanodomains in the plasma membrane of T cells

On both Jurkat and primary T cells, and independent of CAR specificity, *d*STORM images indicate that CARs are organized as monomers and in nanodomains in the cell membrane (Fig. 1a-b, Extended Data Fig. 1 and Supplementary Fig. 1-3). In order to exclude that antibody-induced cross-linking of CARs caused the formation of CAR nanodomains^36^, Jurkat cells expressing a CD19-specific CAR were stained using different antibody incubation times, fixed and then imaged. If antibody labeling causes cross-linking of CARs, we would expect to see an increasing number of nanodomains with increasing incubation time. Strikingly, CAR nanodomains were visible already after 10 min of antibody staining and their size did not change with antibody incubation time suggesting that CAR nanodomains are real and not induced by antibody crosslinking (Extended Data Fig. 2).

To study the distribution of plasma membrane CARs in more detail, we used a density-based spatial clustering of applications with noise (DBSCAN) algorithm with customized localization analysis (LOCAN)^37^. After selecting basal membrane regions that do not show folded membrane areas, localization clusters (≥ 3 localizations per cluster) originating from one or more spatially inseparable primary AF647-labeled antibodies were identified using the DBSCAN algorithm (minPoints = 3, epsilon = 20 nm). The analysis parameters were selected by checking the clustering approach on simulated data^38^. The overall density of CARs in the plasma membrane was low enough to yield well separated nearest neighbors for all plasma membranes investigated. In each experiment, more than 15 cells were analyzed to obtain CAR density distributions. Under the applied experimental conditions, we localized individual antibodies labeled with AF647 at a DOL of 3.0 on average 10 ± 2 times. Hence, absolute localization numbers can be divided by 10 to estimate the number of CARs present on the plasma membrane.

We compared the distribution of localizations per cluster and convex hull areas of localization clusters detected for CD19-specific CAR-T cells and untransduced control T cells (Figure 1d,e). The histograms clearly show that localization clusters analyzed from CAR-T cells contain more localizations and are larger in area as compared to control T cells indicating the co-localization of several CARs in nanodomains. As an additional reference, we compared the number of localizations detected per localization cluster for Jurkat cells stained for the monomeric T cell co-receptor CD45 involved in TCR signaling (Supplementary Fig. 4)^39^. Moreover, we plotted the cumulative histograms of localizations per cluster weighted by the localizations per cluster (Fig. 1f). The plot reflects the percentage of CD19-specific CAR labels found within localization clusters with increasing localizations per cluster and confirms the presence of nanodomains on the plasma membrane of CD19-specific CAR-T cells. Assuming that localization clusters of more than 40 localizations represent CAR nanodomains, we determined the median density of monomers and nanodomains to (3.04 ± 0.9) and (0.69 ± 0.4) μm^-2^, respectively (Fig. 1g), whereby CAR nanodomains contain on average ∼20 CARs. Importantly, this value is a lower bound estimation of the amount of CARs per nanodomain, because steric hindrance will impede quantitative labeling of all CAR epitopes in densely packed protein nanodomains by IgG antibodies.

#### CAR nanodomain formation is independent of CAR expression level and gene transfer method

Given the presence of CAR nanodomains in the plasma membrane, we investigated which potential factors can influence their formation, including restrictions by mechanisms involved in shuttling of the receptors to the plasma membrane, as the natural TCR complex is subject to tightly regulated complex assembly during shuttling. Other potential influences may be intrinsic to the design of CAR-binders or structural domains or signaling-dependent. Furthermore, nanodomain formation can occur signaling-independent mediated through intracellular domains of the receptors^40^. Most prominently, nanodomain formation could be promoted simply by increased abundancy of receptors on the cell surface.

To investigate whether the formation of CAR nanodomains depends on the expression level, we plotted the percentage of CAR monomers and nanodomains as a function of the overall CAR expression level (Extended Data Fig. 3a). These data clearly show that the majority of CARs is organized in nanodomains even at very low overall CAR expression. Hence, our data confirm that ∼20% of all localization clusters (fluorescence signals) detected represent CAR nanodomains that contain ∼68% of all CARs expressed whereas the majority of fluorescence signals (∼80%) can be attributed to monomeric CARs corresponding to ∼32% of all CD19-specific CARs expressed on the cell surface. Furthermore, we checked whether the cell size has an impact on CAR density. Therefore, we calculated and plotted the area of the basal membrane for different conditions and found no significant difference (Extended Data Fig. 3b). Next, we plotted the areas versus the inverse of the localization density and found no correlation between CAR localization density and cell size for all conditions (p-values 0.53 and 0.40 for LV and SB, respectively) (Extended Data Fig. 3c).

To determine the influence of the gene transfer method on CAR nanodomain formation, we investigated the distribution of CD19-specific CARs on CD4^+^ and CD8^+^ T cells using two different Sleeping Beauty (SB)-transposon vectors: a conventional transposon vector, and a minimal expression vector termed minicircle (MC) transposon^41^. Cluster analysis of *d*STORM images showed only small differences between CD4^+^ and CD8^+^ T cells and revealed that the nanodomain formation tendency is similar for the two classes of transposon vectors (Fig. 2a-c and Supplementary Fig. 5). Overall, these results confirm that 58-70% of all CD19-specific CARs expressed are organized in CAR nanodomains on the cell surface of T cells independent of the introduction method (Fig. 2b,c). Thus, it appears that the formation of CAR nanodomains in the plasma membrane represents an intrinsic feature of CAR-T cells.

**Fig. 2.**
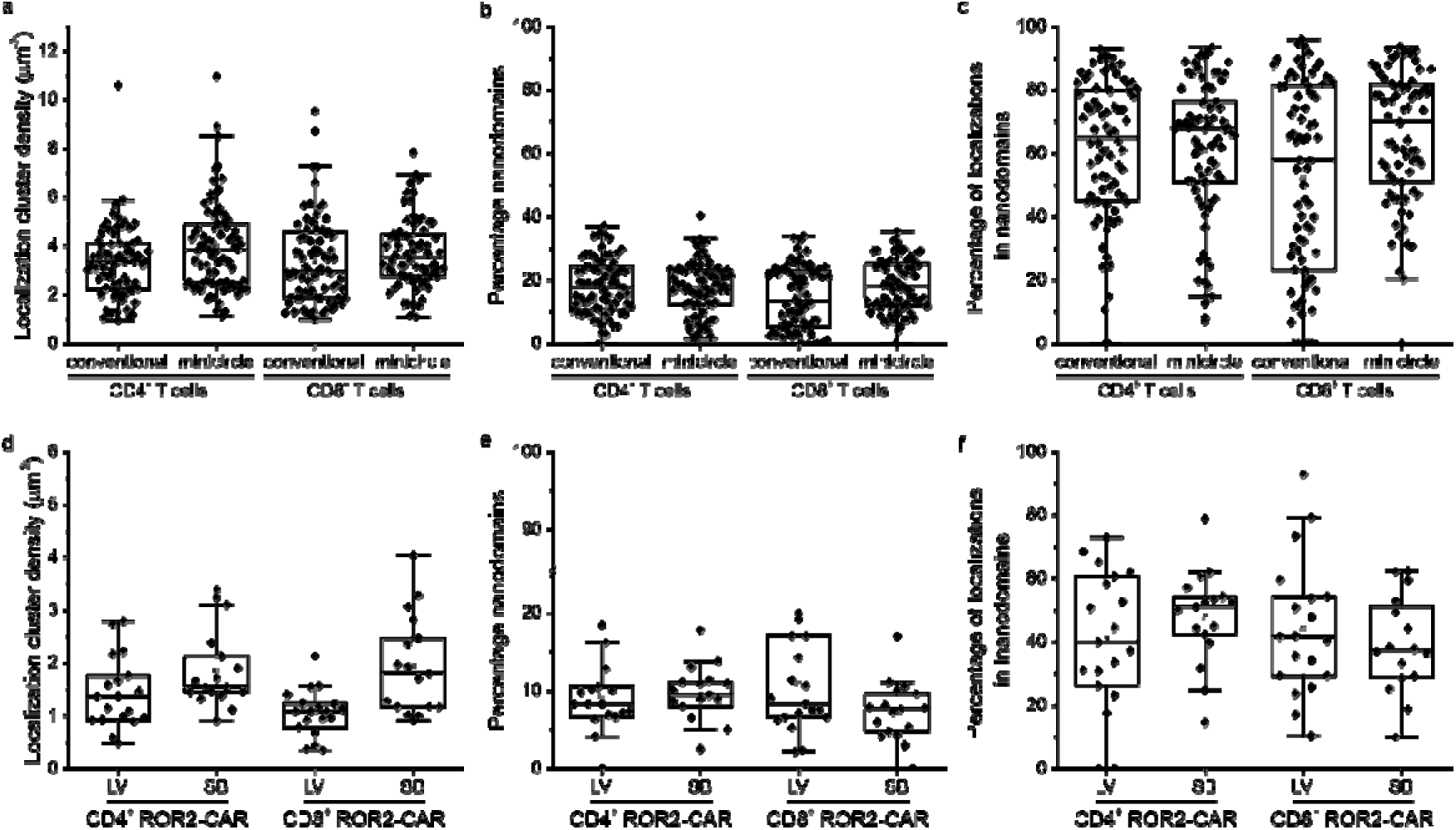
Impact of gene-transfer method on CAR expression and nanodomain formation efficiency on T cell surfaces. **a,** CAR expression (localization cluster densities) of CD19-specific CARs introduced into CD4^+^ and CD8^+^ T cells with conventional SB-transposon plasmids or SB minicircle. **b**, Percentage of large CAR localization clusters detected for different vectors, T cells, and introduction method showing that 13-18% of all localization clusters are CAR nanodomains. **c**, Percentage of CAR localizations found in nanodomains for different T cells and vectors showing that 58-70% of all CARs expressed on the plasma membrane are organized in nanodomains. **d,** CAR expression (localization cluster densities) of ROR2-specfic CARs introduced into CD4^+^ and CD8^+^ T cells by lentiviral transduction (LV) and SB MC transfection. **e,** Percentage of large ROR2-specific CAR localization clusters detected showing that 8-9% of all localization clusters are CAR nanodomains. **f**, Percentage of ROR2-specific CAR localizations found in nanodomains using different introduction methods (LV and SB) showing that 38-51% of all ROR2-specific CARs expressed on the plasma membrane are organized in nanodomains.

Finally, we tested whether distinct gene transfer methods, namely lentiviral transduction and SB MC nucleofection, influence CAR expression and nanodomain formation of CARs in the plasma membrane, and how this translates to CAR-T cell functionality. Even though the CAR expression level was not distinguishable by flow-cytometry, *d*STORM revealed substantially lower ROR2-specific CAR localization cluster densities on CD4^+^ T cells of (1.37 ± 0.4) µm^-2^ for LV transduction than for SB transfection (1.55 ± 0.2) µm^-2^ (Fig. 2d and Extended Data Fig. 4). LV and SB-mediated gene transfer leads to distinct patterns of genomic insertions^41^, which translate to distinct spatiotemporal gene expression patterns and CARs expressed in the plasma membrane. Using LV and SB-mediated gene transfer, we found that only 8-9% of all localization clusters represent nanodomains that contain 38-51% of all ROR2-CARs expressed (Fig. 2e,f).

### Gene transfer method and expression level influence CAR-T cell function

To verify the functionality of ROR2- and CD19-specific CAR-T cells we investigated their cytotoxic capacity. Here, an advantage of SB-generated ROR2-specific CAR-T cells over LV-generated ROR2-specific CAR-T cells was apparent, with 3 out of 4 donors displaying significant advantages in short-term cytotoxicity (5 h) for ROR2-specific CAR-T cells and in 2 out of 3 donors for CD19-specific CAR-T cells (Fig. 3a,c, Extended Data Fig 5). At higher effector-to-target ratios (E:Ts), SB ROR2-specific CAR-T cells exhibited on average 18-30 % of short-term specific lysis against ROR2^+^ MDA-MB-231 cells, while LV ROR2-specific CAR-T cells showed 14-23 % of specific lysis (Fig. 3b). SB CD19-specific CAR-T cells were able to achieve 32-48 % of short-term specific lysis on average at higher E:Ts, and LV CD19-specific CAR-T cells achieved 29-42% (Fig. 3d). However, LV CAR-T cells of both specificities were able to compensate for the initial lag-phase during long-term follow-up (≥ 20 h), equalizing the final anti-tumor potency in this assay with an average specific lysis of 82-92% for SB ROR2-specific CAR-T cells, 78-89% for LV ROR2-specific CAR-T cells, and for both SB and LV CD19-specific CAR-T cells (Fig. 3b,d). The trend of functional advantage of SB-generated CAR-T cells regardless of specificity was also observed in cytokine secretion assays, with ROR2-specific SB CAR-T cells secreting significantly higher amounts of IFNγ and slightly but not significantly higher amounts of IL-2 (Fig. 3g,h), while no significant differences between the two gene transfer methods were observed in terms of proliferation capacity (Fig. 3e,f).

**Fig. 3.**
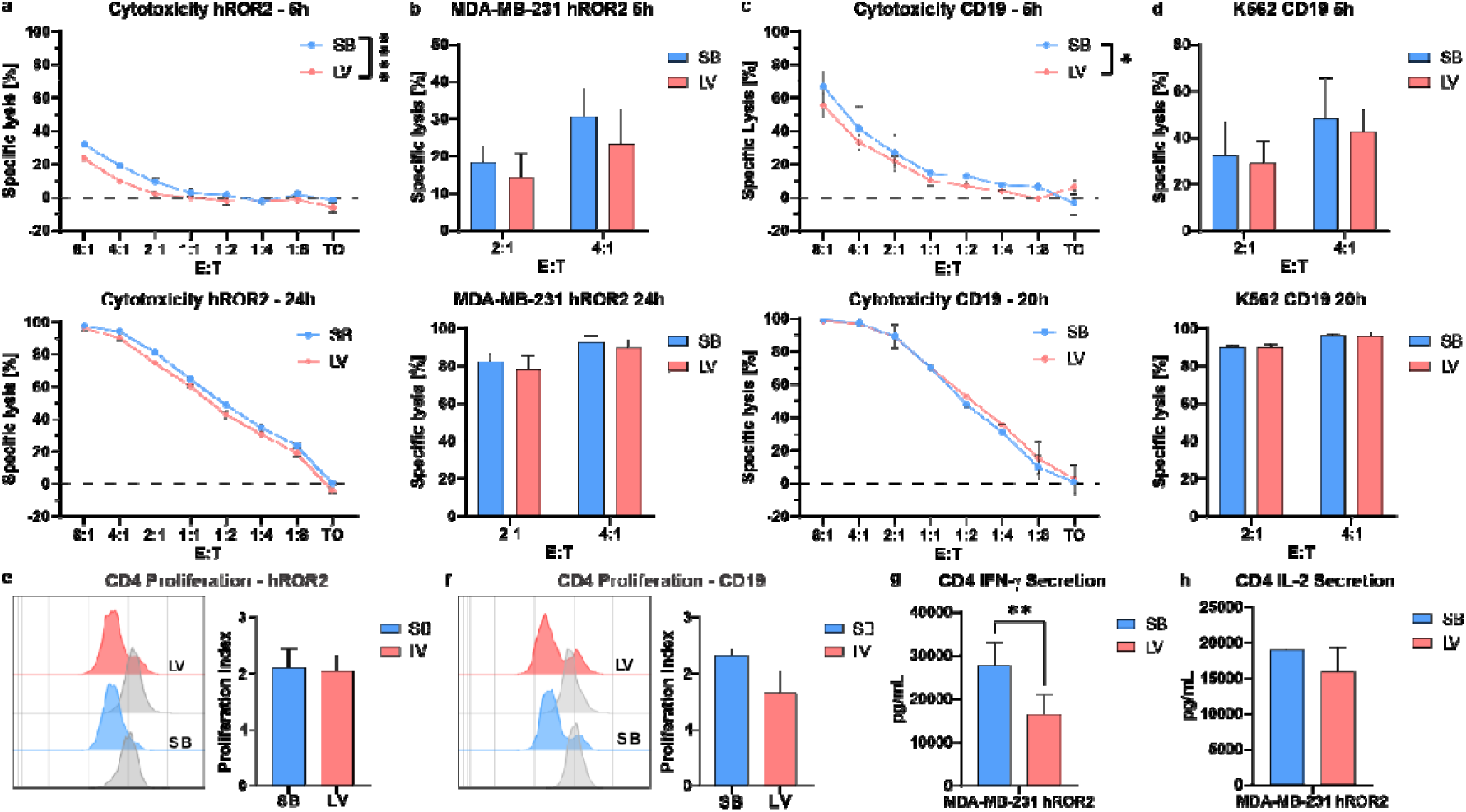
SB CAR-T cells show tendencies of functional advantage compared to LV CAR-T cells, independent from nanodomain formation. Functional *in vitro* characterization of CAR-T cells generated by Sleeping Beauty transposition (SB) or lentiviral transduction (LV), targeting either hROR2 or hCD19. **a-d**, short (5h) or long-term (≥ 20h) antigen-specific lysis mediated by CD8^+^ CAR-T cells co-cultured with MDA-MB-231 (hROR2^+^) or K562 (CD19^+^) in multiple effector-to-target ratios (E:T) in luminescence-based cytotoxicity assays. **a**,**c**, Cytotoxicity of one donor representative for 4 (hROR2) or 3 (CD19) biological replicates. Data are depicted as Mean ± SD of 3 technical replicates. Statistical analyses were performed using two-way ANOVA with Sidak correction for multiple comparisons. **b**,**d**, Average cytotoxicity of 4 (hROR2) or 3 (CD19) donors at E:Ts of 4:1 or 2:1. Data are represented as mean ± SEM. **e**,**f**, proliferation of CD4^+^ CAR-T cells co-cultured with target-positive tumor cells for 72h. Data shows representative proliferation peaks of CFSE-labelled CAR-T cells (colored) or control T cells (grey) and average ± SEM proliferation index (PI) of 5 (hROR2) or 3 (CD19) biological replicates. **g**,**h**, Cytokine secretion of CD4^+^ CAR-T cells incubated with target-positive tumor cells for 24h. Data are represented as mean ± SEM of 5 biological replicates. Statistical analysis was performed using two-tailed ratio-paired Students T-test. *p< 0.05, **p< 0.01, ***p< 0.001, ****p< 0.0001.

Thus, regardless of the unaltered CAR distribution, a context dependent functional advantage of SB-generated CAR-T cells over LV-generated CAR-T cells was observed. This may be attributed to the slight increase in CAR surface expression that was invisible to flow cytometry but could be visualized by *d*STORM.

### CAR design does not affect nanodomain formation

Since expression levels and the transduction method did not have an effect on CAR nanodomain formation efficiency, we investigated whether modifications in the spacer or linker domain of the construct lead to changes in CAR distribution. To this end, four complementary CAR constructs with CD19 specificity were generated (Fig. 4a). For one pair, the linker connecting scFv heavy and light chains was shortened by removing the upper hinge (UH) sequence and the scFv was combined with either the previously used IgG spacer and CD28-based transmembrane domain (SL-CD28tm) or an alternative CD8α-based spacer and transmembrane domain (SL-CD8α), respectively. For the second pair, the original scFv-Linker was combined with the two spacer/transmembrane variants (LL-CD28tm; LL-CD8α). Although the expression levels and surface densities of CAR constructs with a CD8α transmembrane domain were lower (Fig. 4b), we did not find significant differences in the percentage of nanodomains (9-12%) or in the percentage of CARs contained in nanodomains (40-52% on average) (Fig. 4b-d and Supplementary Fig. 6). Upon functional characterization of the distinct CAR designs we did not observe differences in short-term cytotoxicity (5 h) against the CD19-positive tumor cell line JeKo-1 between any of the constructs, showing indistinguishable rates of specific cytotoxicity throughout different E:Ts (Fig. 4e). Interestingly, we did observe differences in short-term cytotoxicity (5 h) against CD19-positive Raji cells revealing a general advantage in specific lysis with 54%, 76%, and 89% for the CD28tm-constructs compared to 41%, 62%, and 82% for the CD8α-variants throughout different E:Ts, respectively (Fig. 4f). These results illustrate that formation of CAR nanodomains is independent of CAR design but CAR surface expression directly influences the functional potency.

**Fig. 4.**
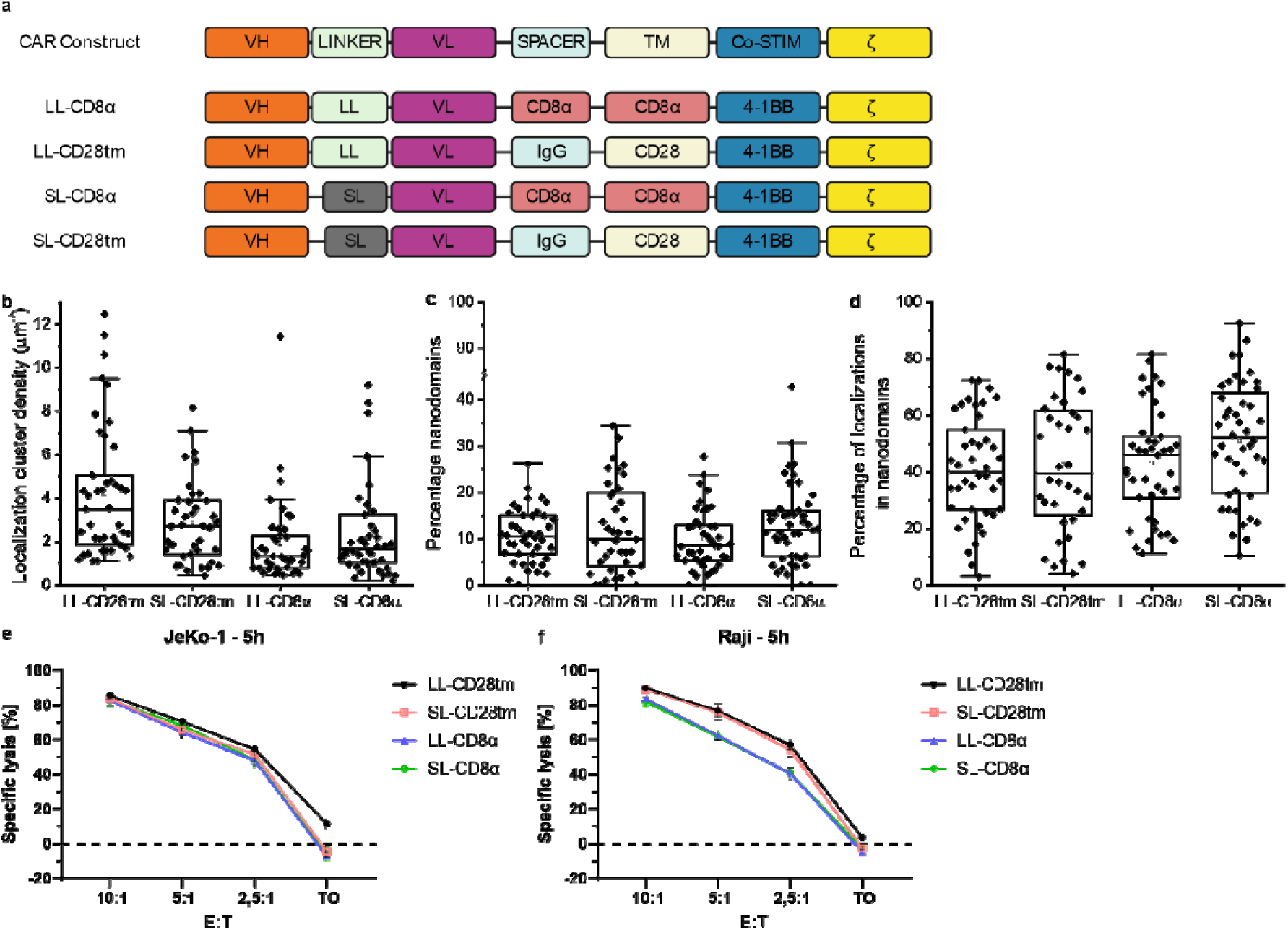
Nanodomain formation is independent of CAR construct design. **a**, schematic depiction of CAR construct designs. VH= variable domain heavy chain; VL = variable domain light chain; LL = longer linker; SL = shorter linker; TM = transmembrane domain. **b**,**c,d,** *d*STORM-based analysis of CAR expression and organization. Boxplots depict the localization cluster density (b), percentage of nanodomains (c) and the percentage of localizations in nanodomains (d) for the different CAR designs. **e**, short-term cytotoxicity assay of CAR-T cells transduced with the distinct CD19-CAR designs against CD19-positive JeKo-1 cell line for multiple E:Ts. Data are represented as Mean ± SEM. **e**, Short-term cytotoxicity assay of CAR-T cells transduced with the distinct CD19-CAR designs against CD19-positive Raji cell line for multiple E:Ts. Data are represented as Mean ± SEM. Statistical analysis was performed by two-way ANOVA with Sidak correction for multiple comparison.

### CAR monomers and nanodomains accumulate at tumor contact sites

To investigate the involvement of CAR monomers and nanodomains in T cell engagement and immunological synapse formation, we transduced CD3 positive CD19-specific CAR-Jurkat cells additionally with GFP tagged ZAP70, a kinase that is recruited to phosphorylated CD3ζ chains upon successful T cell activation^42, 43^. ZAP70 acts as a critical effector of downstream signaling after the initial engagement of the TCR and accumulates at the immunological synapse. K562 cells that were stably transfected to overexpress CD19 and CD20 (K562_CD19^+^_CD20^+^) were co-cultured with CD19-specific CAR-Jurkat cells at a ratio of 1:10 for 30 min and then immunostained with an anti-CD20 antibody for visualization of tumor cells before imaging. Reference experiments were performed with ZAP70-GFP expressing Jurkat cells that were not equipped with CD19-specific CARs. Fluorescence immunolabeling of CD20 on the tumor cell surface was preferred over CD19 to enable recognition of the antigen by CD19-specific CARs. While Jurkat cells without CARs showed a homogeneous distribution of the ZAP70-GFP signal (Extended Data Fig. 6a-c), it accumulated at contact sites to tumor cells in CD19-specific CAR-Jurkat cells. Interestingly, ZAP70 also accumulated at tumor cell cytoplasmic projections (filopodia) embracing the T cells (Extended Data Fig. 6d-f).

Next, we investigated if CARs also accumulate at contact sites of tumor and CD19-CAR equipped Jurkat cells. Therefore, we performed three-color imaging experiments with ZAP70-GFP expressing CD19-specific CAR-Jurkat cells. CD20 on tumor cells and CARs on T cells were labeled with primary antibodies. Fluorescence images clearly displayed that both, the ZAP70 and CAR signal accumulate at tumor cell contact sites (Fig. 5a-d). Control experiments performed in the absence of tumor cells showed that ZAP70 and CAR fluorescence signals are homogeneously distributed over the entire plasma membrane of CAR-Jurkat cells (Supplementary Fig. 7). In addition, we performed identical experiments with primary CAR-T and tumor cells. The obtained results confirm that ZAP70-GFP accumulates at contact sites in CD4^+^ and CD8^+^ T cells expressing both ZAP70-GFP and CD19-CAR (Supplementary Fig. 8a-c and 8g-i) but not at contact sites between tumor cells and ZAP70-GFP expressing primary cells that are not equipped with CD19-CARs (Supplementary Fig. 8d-f and 8j-l).

**Fig. 5.**
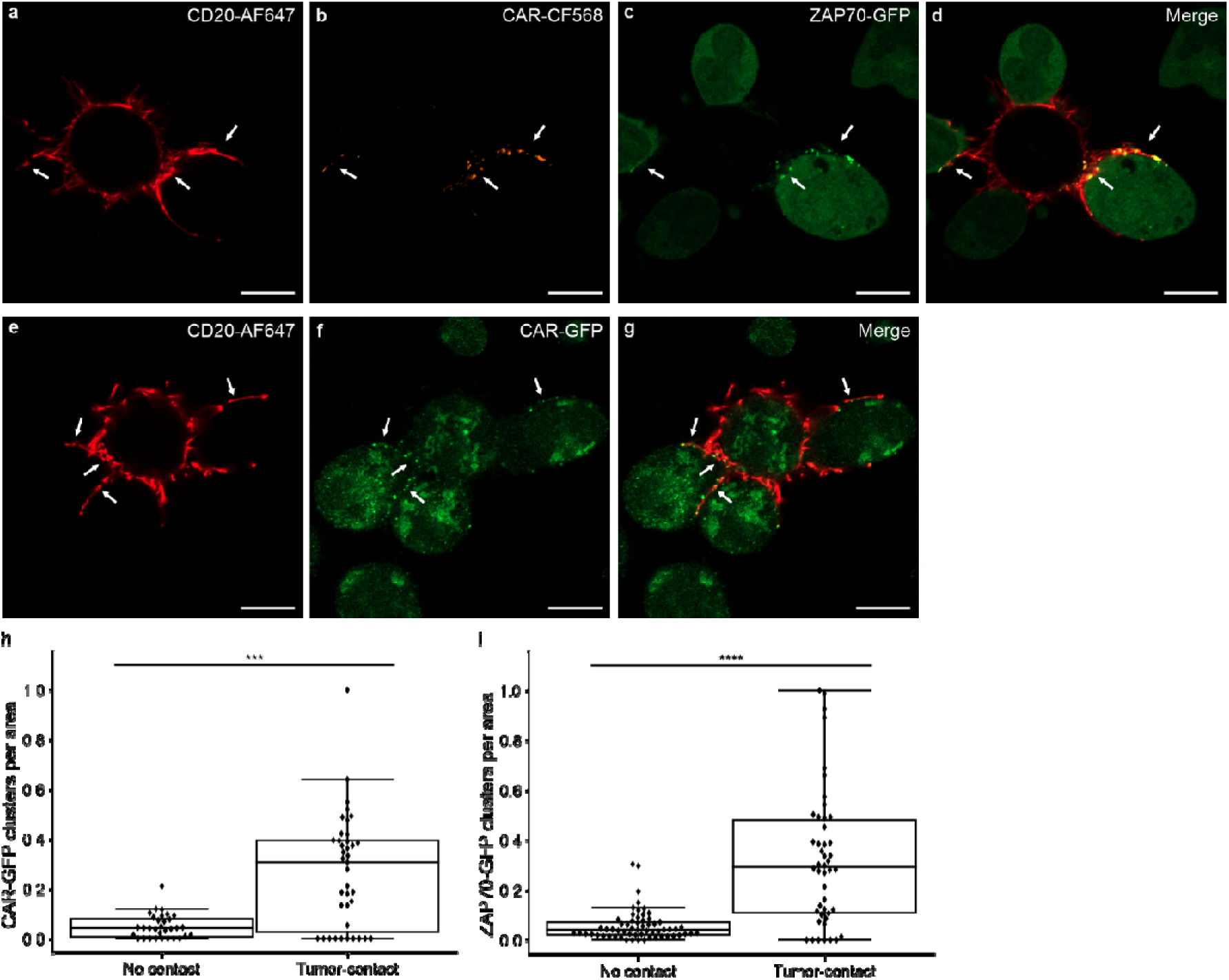
CAR and ZAP70 accumulate at tumor contact areas. **a-c**, Airyscan confocal fluorescence images of K562_CD19^+^_CD20^+^ tumor cells co-cultured with CD19-specific CAR-Jurkat cells expressing ZAP70-GFP for 30 min before staining with an anti-CD20 antibody labeled with AF647 and fixation (a), anti-CD19-CAR antibody labeled with CF568 (b), and the GFP signal (c). **d**, Merged image showing the accumulation of CAR and ZAP70 at tumor contact sites. **e,f**, Airyscan confocal fluorescence images of K562_CD19^+^_CD20^+^ tumor cells co-cultured with CD19-specific CAR-GFP Jurkat cells for 30 min before fixation and staining with an anti-CD20 antibody labeled with AF647 (e). The GFP signal of the CAR is shown in (f). **g**, Merged image showing the accumulation of CARs at tumor contact sites. **h,** Quantification of GFP-signal for CD19-CAR-GFP expressing Jurkat cells. The GFP signal in airyscan images (maximum z-projection) at cell-cell contact areas and the number of accumulations counted. Tumor-cell contact areas were identified by the presence of CD20-AF647 staining. On average 0.05 and 0.27 CAR-GFP clusters per area were detected without and with tumor contact, respectively. **i,** Quantification of GFP-signal for ZAP70-GFP expressing Jurkat cells. The GFP signal in airyscan images (maximum z-projection) at cell-cell contact sites and the number of accumulations counted. Tumor-cell contact sites were identified by the presence of CD20-AF647 staining. On average, 0.06 and 0.32 ZAP70-GFP clusters per area were detected without and with tumor contact, respectively. White arrows indicate contact sites. Scale bars, 5 µm.

To rule out antibody induced clustering of CARs at contact sites to tumor cells, we performed additional experiments with Jurkat cells transfected with a CD19-specific CAR tagged with GFP. Two-color fluorescence images supported our finding that CARs move towards the contact sites to tumor cells. (Fig. 5e-g). Closer inspection of CAR distributions on CD19-specific cells in the absence (Supplementary Fig. 7) and presence of tumor cells (Fig. 5b,f) demonstrates unequivocally that CAR nanodomains accumulate at tumor contact sites. To quantify the accumulation of CAR and ZAP70 molecules at tumor contact sites we analyzed the GFP signals measured at contact sites between Jurkat and tumor cells. Since confocal airyscan fluorescence imaging does not exhibit exquisite single-molecule sensitivity this analysis captures mainly nanodomains of ZAP70 and CAR molecules, respectively. Thus, our data clearly indicate that CAR nanodomains accumulate at tumor contact sites and recruit ZAP70 (Fig. 5h,i). After 30 min of co-culture, we found on average 0.27 CAR nanodomains µm^-2^ and 0.32 ZAP70 nanodomains µm^-2^ at tumor contact sites (Supplementary Fig. 9).

To investigate if CAR monomers move towards and accumulate at tumor contact sites as well, we performed correlative Structured Illumination Microscopy (SIM) and *d*STORM experiments with anti-CAR antibodies in the absence and presence of tumor contacts. These data unequivocally confirm that the cell membrane is depleted of CAR monomers and nanodomains after contact formation with a tumor cell. This finding indicates that both, CAR nanodomains and monomers accumulate at contact sites to tumor cells (Extended Data Fig. 7).

### CAR-T cells form multifocal immunological synapses

CARs have been shown to employ the TCR-CD3ζ downstream signaling axis and have been suggested to form immunological synapses analogous to the classical TCR “bull’s eye” that is both, central to effector function as well as inter- or intracellular communication and has recently been linked to clinical outcome of CAR-T cell immunotherapy^9,29–32^. To gain a more detailed understanding of the molecular interactions during immunological synapse formation we utilized live-cell fluorescence imaging to study the *real time* interaction between ZAP70-GFP expressing CAR-T cells and anti-CD20 ATTO643-stained Raji cells. In particular, we used lattice light-sheet microscopy (LLSM)^44^ for whole-cell volumetric imaging of immunological synapse formation between CAR-T and tumor cells at high temporal resolution. Since out-of-focus regions are not illuminated, phototoxicity is drastically reduced and enables long-term observations of interacting cells. Three-dimensional, whole-cell, live, and gentle imaging with LLSM also facilitates a major step forward in designing experimental systems for immunological synapse imaging as compared to earlier strategies which mainly used horizontally orientated synapses between immune cells and artificial substrates mimicking target cells, such as antibody-coated glass slides or supported planar lipid bilayers with anchored ligand proteins as surrogates for APCs^21, 28, 32, 45^. While these strategies enabled the application of super-resolution microscopy techniques^46^ still preserving the essential hallmarks of T cell signaling, it is likely that such artificial substrates do not recapitulate many of the complexities in intercellular interactions.

To visualize immunological synapse formation by LLSM, we used a monolayer of Raji cells immunostained with ATTO643 labeled anti-CD20 antibodies and added primary CD4^+^ and CD8^+^ CD19-specific CAR-T cells expressing ZAP70-GFP prior to start of the experiment. Two-color fluorescence images were acquired at 37°C and 5% CO_2_ every minute for a total duration of 40 min. First, we determined the duration necessary for CAR-T and Raji cells to take up contact. Here, we measured an average contact formation time of (13.5 ± 3.3) min and (12.7 ± 1.5) min (averaged over 5 different contact events in each case) for CD4^+^ and CD8^+^ CAR-T cells respectively, after they were added to the monolayer of Raji cells (Supplementary Movies S1-S4). Interestingly enough, these contacts remained stable during the entire acquisition period of 40 min. Upon contact formation, ZAP70 molecules accumulated at contact sites to tumor cells and formed multiple ZAP70 clusters, which is indicated by higher GFP fluorescence intensity regions in contact zones (Fig. 6a-b and Supplementary Fig. 10a) similar to our observations by confocal microscopy (Fig. 5 and Extended Data Fig. 5,6).

**Fig. 6.**
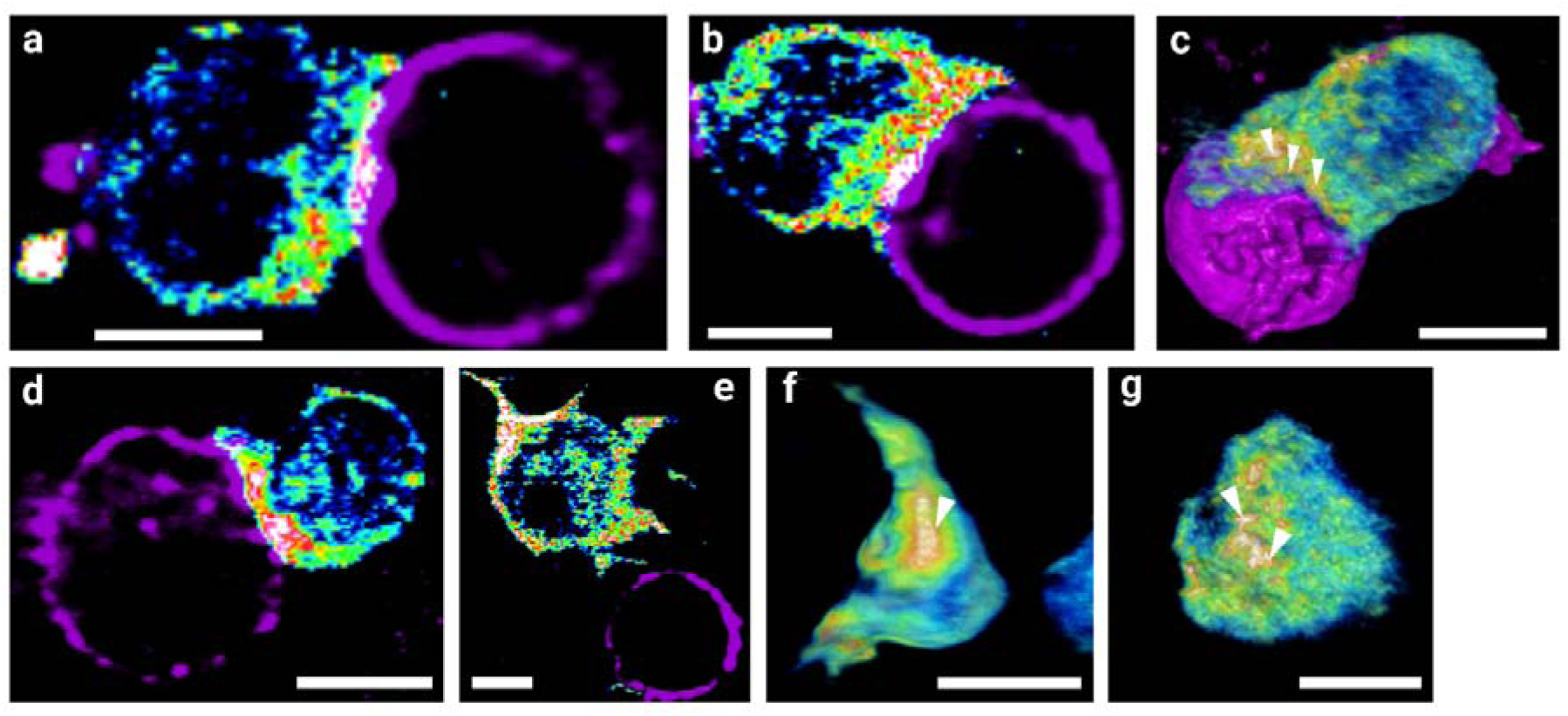
Lattice light-sheet imaging of immunological synapse formation. **a,** Immunological synapse formation between a representative CD4^+^ T cell expressing CD19-specific CARs and ZAP70 labeled with GFP (rainbow LUT), and a Raji cell (magenta) stained for CD20 by antibodies conjugated to ATTO643. **b,** A second example of synapse formation between a CD4^+^ CD19-specific CAR-T cell and a Raji cell. An increase in fluorescence signal of GFP channel (rainbow LUT) can be seen at the cell-cell junction confirming ZAP70 accumulation at the cell-cell contact site. **c**, 3D volume rendering of a CD4^+^ CD19-specific CAR-T cell and a Raji cell forming an immunological synapse. Scattered accumulation of GFP signal (white pointers) can be visualized at the contact site. **d,** Immunological synapse formation between a representative CD4^+^ T cell (rainbow color scale) without CAR construct expressing ZAP70-GFP and a Raji cell (magenta). Synapse formation was induced by addition of staphylococcal enterotoxin type E (SEE). **e,** No immunological synapse formation is seen between CD4^+^ T cells without CAR construct expressing ZAP70-GFP and Raji cells. **f,** Side view of a CD4^+^T cell without CD19-CAR at the site of synapse formation (induced by addition of SEE) shows accumulation of ZAP70-GFP signal resembling the classical ‘bull’s eye’ organization. **g,** Scattered clustering of ZAP70-GFP at multiple foci (indicated with white arrows) interleaved by lower fluorescence intensity regions unlike the classical organization can be seen at the site of synapse formation for a CD4^+^ CD19-specific CAR-T cell (right panel). Scale bars, 5 µm.

Fortunately, the whole-cell volumetric imaging capability of LLSM enabled us to visualize the contact zone in more detail and reveal the formation of multifocal immunological synapses between CAR-T and tumor cells (Fig. 6c,g, Supplementary Fig. 10b). We also performed reference experiments with primary ZAP70-GFP T cells that do not express CARs. These cells, showed contact formation after (14.2 ± 1.7) min of addition of T cells but the contacts remained stable only for ∼0.8 min (Supplementary Movie S5 and S6), without any indication of ZAP70-GFP accumulation at contact sites (Fig. 6e, Supplementary Fig. 10c,d). This suggests that the multifocal immunological synapse observed, is indeed induced by the CARs.

Furthermore, we performed identical experiments with primary CD19-specific CD4^+^ and CD8^+^ CAR-T cells and donor-matched primary B cells. Here too, contacts with B cells were formed within (14.1 ± 2.4) and (15.5 ± 2.7) min (Supplementary Movie S7 and S8) and the ZAP70-GFP signal accumulated at tumor contact sites (Supplementary Fig. 11a,b and Supplementary Fig. 12a). In contrast, when T cells were not equipped with a CAR, no or only transient contacts were established between T and B cells (Supplementary Fig. 11c and 12b). Finally, we validated whether a conventional “bull’s eye”-shaped immunological synapse is formed by primary T cells expressing ZAP70-GFP by pulsing Raji cells with staphylococcal enterotoxin type E (SEE). Here we determined an average contact formation time of (15.1 ± 2.6) min for CD4^+^ and (17.3 ± 1.5) min for CD8^+^ cells (Supplementary Movies S9 and S10) similar to the time span determined in our experiments with CD19-specific CAR-T cells. However, here the ZAP70-GFP signal accumulated at the contact sites to tumor cells and 3D analysis of lattice light-sheet data revealed the formation of a typical “bull’s eye”-shaped immunological synapse with distinct peripheral and central supramolecular activation complex areas (pSMAC and cSMAC) indicated by TCR-mediated accumulation of ZAP70-GFP (Fig. 6d,f, Supplementary Fig. 10c).

## Discussion and Conclusion

The improved understanding of cancer immunobiology, and the leverage of this knowledge to effectively eradicate malignant cells has revolutionized the field of cancer therapeutics. In particular, genetically modified CAR-T cells have been a breakthrough because of their successful use for the treatment of advanced leukemias and lymphomas including cure of previously incurable diseases^4^. The constantly growing number of CAR-T cell targets emphasizes the potential for improved future immunotherapies. On the other hand, CAR-T cell therapy exhibits limitations that must be addressed in order to translate its full potential also to the treatment of solid tumors and to reduce side effects in hematological malignancies^7, 47^. Despite the very favorable clinical responses, we are currently still far away from a detailed mechanistic picture of the molecular mode of function of CAR-T cells. Here, methods that enable the visualization of CAR expression, distribution and interaction with tumor cells with molecular resolution are required to further improve the efficiency and safety of CAR-T cell immunotherapies.

Therefore, we used single-molecule sensitive super-resolution fluorescence imaging to determine the expression and distribution of CARs on the plasma membrane of primary T and Jurkat cells. *d*STORM images demonstrated that the majority of CARs (up to ∼70%) is organized in nanodomains on the surface of Jurkat and primary T cells (Fig. 1 and Extended Data Fig. 1-3). This finding was independent of T cell size, CAR specificity and introduction method. Only for ROR2-specific CARs using lentiviral transduction (LV) and SB MC nucleofection, T cells showed lower overall expression levels whereby 38-51% of all CARs are still organized in nanodomains in the plasma membrane (Fig. 2). Since we used fluorescently labeled primary antibodies in our CAR quantification experiments, the measured expression levels represent lower bounds because of the limited antigen accessibility on the basal plasma membrane of adherent cells. However, 3D lattice light-sheet *d*STORM experiments have also shown that receptor distribution and mobility are largely unaffected by contact to the coverslip^48^.

Functional characterization of LV or SB-engineered ROR2 and CD19 specific CAR-T cells revealed functional advantages for SB CAR-T cells in short-term cytotoxicity assays, regardless of CAR specificity (Fig. 3) that could not necessarily be attributed to differences in nanodomain formation and did not persist in long-term cytotoxicity assays (≥ 20 h). SB ROR2 CAR-T cells also showed a significant advantage in IFNγ production and a similar advantageous trend was observed for IL-2 secretion independent of CAR-T cell specificity. In accordance with long-term cytotoxicity data, the advantage of SB CAR-T cells was negligible in proliferation assays, potentially due to masking of a head start of SB CAR-T cells/a lag-phase of LV CAR-T cells in long-term integrative or end-point analyses, respectively. Here, *d*STORM clearly resolved differences in localization cluster densities between the gene transfer methods that were not detectable by flow cytometry. Strong differences in CAR expression, visible even by flow cytometry, were recently shown to correlate with clinical outcome in BCMA CAR-T cell therapy prompting the attribution of the observed functional effects to distinct CAR expression levels rather than other putative factors^49^.

Furthermore, comparison of distinct linker, spacer and transmembrane configurations in CAR design suggest that the formation of CAR nanodomains is independent of CAR construct design (Fig. 4). Interestingly, the incorporation of a CD8α-based spacer and transmembrane domain resulted in decreased short-term cytotoxic activity against Raji cells (Fig. 4). Again, these results point to the distinct localization cluster densities found for constructs with the CD8α-based spacer and transmembrane domain. Thus, our data suggests a link between CAR surface expression and enhanced short-term cytotoxicity as well as cytokine production.

In addition, we employed confocal microscopy on fixed and lattice light-sheet microscopy on living cells for the visualization of tumor-T cell interactions. Intriguingly, these experiments demonstrated that CAR monomers and nanodomains as well as the downstream effector molecule ZAP70 accumulate at contact sites in multiple foci (Fig. 5, Extended Data Fig. 6,7 and Supplementary Fig. 8,9). For conventional T cell engagement, we were able to image the well described classical bull’s eye shape of the immunological synapse by pulsing tumor cells with SEE and co-incubation with primary T cells that express a GFP-tagged ZAP70 (Fig. 6). In contrast, all CAR-T cells formed multifocal immunological synapses with tumor cells (Fig. 6). Interestingly, we were also able to capture the interactions between CD19-specific CAR-T cells and donor-matched primary B cells (Supplementary Fig. 12). B cell aplasia, caused by the killing of B cells by CD19-CAR-T cells, is a very common on-target off-tumor adverse effect in patients treated with CD19-specific CAR-T cells. Hence LLS imaging could be used advantageously to map on-target off-tumor effects, even when the antigen expression is too low for flow cytometry detection^3^.

## Acknowledgements

The authors thank E. Maier, B. Grams and L. Winkelmann for support with cell culture and cloning. The project has been funded by the *Bundesministerium für Bildung und Forschung* (BMBF, grant #13N15986, “IMAGINE”) and the *Deutsche Forschungsgemeinschaft* (DFG, German Research Foundation) - SFB/TR 338 “LETSimmun” - A05/A02. M.S and S.R. received funding from the European Research Council (ERC) under the European Union’s Horizon 2020 research and innovation programme (grant agreement No 835102). MH received funding from the Innovative Medicines Initiative 2 Joint Undertaking under grant agreement Nos 116026 and 853988. These Joint Undertakings receive support from the European Union’s Horizon 2020 research and innovation program, EFPIA and JDRF INTERNATIONAL.

## Author contributions

T.N., M.H., H.E., and M.S. conceived and designed the project. T.N., M.H. and M.S. supervised the project. C.V. performed all *d*STORM and confocal airyscan microscopy experiments. L.G. and J.W. generated CAR constructs. L.G. generated CAR-T cells and performed quality control and evaluation of CAR-T cell effector function. C.V. and N.B. performed SIM/*d*STORM experiments. C.V. and A.G. performed lattice light-sheet microscopy experiments. C.V., S.R. and S.D. performed data analysis. T.N., M.H. and M.S. wrote the manuscript. All authors revised the final manuscript.

## Data availability

The data that support the findings of this study will be provided by the corresponding author upon reasonable request. The code used for super-resolution microscopy analysis is available on GitHub (https://github.com/super-resolution/Verbruggen-et-al-2023-supplement).

## Methods

### Human subjects

Blood samples were obtained from healthy donors who provided written informed consent to participate in research protocols approved by the Institutional Review Board of the University of Würzburg. Peripheral blood mononuclear cells (PBMCs) were isolated by density gradient centrifugation using Ficoll-Hypaque (Sigma-Aldrich, St Louis, MO, USA).

### CAR construct generation in SB or LV vectors

The construction of epHIV7 lentiviral vectors containing CARs with 4-1BB costimulatory domain has been described in detail elsewhere^50^. scFvs targeting CD19 (FMC63), ROR1 (R11), ROR2 (4-1) (heavy and light chains connected by human IgG3 upper+middle Hinge) were synthesized by GeneArt (ThermoFisher, Regensburg, Germany) and subcloned into lentiviral backbones containing IgG3- or CD8α-derived spacer domains, a CD8α-derived or a CD28-derived transmembrane domain and an intracellular signaling domain composed of 4-1BB and CD3ζ. All vectors contained a truncated epidermal growth factor receptor (EGFRt) encoded in the transgene cassette downstream of the CAR^51, 52^. The CAR and EGFRt transgenes were separated by a T2A ribosomal skip element. LV constructs were used as basis for the generation of SB transposon plasmid or minicircle vectors as previously described^53^.

#### Preparation of lentivirus

Lentivirus supernatants were produced in 293T cells co-transfected with the respective lentiviral vector plasmids and the packaging vectors pCHGP-2, pCMV-Rev2, and pCMV-G using Calphos transfection reagent (Clontech, Takara, Jusatsu, Japan). Medium was changed 16 hours after transfection, and lentivirus collected after 72 hours. To collect virus particles, ultracentrifugation was performed at 24,900 rpm for 2 hours at 4 °C. Jurkat cells were transduced with increasing amounts of virus for lentivirus titration.

#### Preparation of LV CAR-T cells

LV CAR-T cells were generated as previously described from CD8^+^ or CD4^+^ bulk T cells purified from PBMCs of healthy donors using negative isolation with immunomagnetic beads (Miltenyi Biotec), activated with anti-CD3/CD28 beads according to the manufacturer’s instructions (Life Technologies, ThermoFisher), and transduced into 5×10^5^ by spinoculation on day 1 of activation using lentiviral supernatant at a multiplicity of infection (MOI) of 3 ^50^.

#### Preparation of SB CAR-T cells

SB CAR-T cells were generated as previously described from CD8^+^ or CD4^+^ bulk T cells purified from PBMCs of healthy donors using negative isolation with immunomagnetic beads (Miltenyi Biotec), activated with anti-CD3/CD28 beads according to the manufacturer’s instructions (Life Technologies, ThermoFisher)^53^. Nucleofection of transposon (plasmid or MC) and transposase (MC) vectors was performed on day 2 of activation into 1×10^6^ T cells using 4D-Nucleofector according to the manufacturer’s instructions (Lonza, Cologne, Germany).

#### T cell culture

CAR-T cells were propagated in RPMI-1640 supplemented with 10% human serum (Bavarian Red Cross, Wiesentheid, Germany), 2mM Glutamax (Gibco, ThermoFisher), 100 U/ml penicillin–streptomycin (Gibco, ThermoFisher) and 50 U/ml IL-2 or 50 U/ml IL-2, IL-15 and IL-7 (Miltenyi Biotec). Trypan blue staining was performed to quantify viable T cells. Prior to functional testing, EGFRt-positive T cells were enriched using biotin-conjugated anti-EGFR mAb and anti-biotin beads (Miltenyi Biotec), and subsequently expanded using a rapid expansion protocol based on CD3-TCR-activation via OKT-3 (Miltenyi Biotec) and irradiated donor-mismatched PBMCs (all CAR-T cells, unless indicated) or irradiated feeder cells expressing CD19 or anti-CAR surface-proteins (LV/SB comparison of CD19 and ROR2 CAR-T cells) for 7-14 days.

#### Evaluation of CAR-T cell cytotoxicity

Target cells stably transduced with ffluc/GFP were incubated in triplicate wells at 1×10^4^ cells/well with effector T cells at different effector to target (E:T) ratios or just tumor cells (TO) as indicated. D-luciferin substrate (Biosynth) was added to the co-culture to a final concentration of 0.15 mg/ml, and the decrease in luminescence signal in wells that contained target cells and T cells was measured using a luminometer (Tecan, Männedorf, Switzerland). Specific lysis was calculated in reference to control T cells that been treated equally.

#### Evaluation of cytokine secretion

5×10^4^ CAR-T cells were plated in triplicate wells with target cells at an E:T ratio of 4:1 (CD19) or 1:1 (ROR2), and secretion of IFNγ and IL-2 were measured by ELISA (ELISA MAX, BioLegend) in supernatant removed after 24-hour incubation.

#### Evaluation of proliferation capacity

CAR-T cells were labeled with 0.2 μM carboxyfluorescein succinimidyl ester (CFSE, Invitrogen, ThermoFisher), washed, and plated in duplicate or triplicate wells with irradiated (80 Gy) target cells at an E:T ratio of 4:1 (CD19) or 1:1 (ROR2). No exogenous cytokines were added to the culture medium. After 72-hour incubation, cells were analyzed by flow cytometry to assess cell division of T cells and Proliferation Index, constituting the total number of divisions divided by the number of cells that went into division, was calculated using built in FlowJo V10.0 proliferation modelling (Treestar, Ashland, OR, USA).

### Statistical analysis of functional data

Data from *in vitro* experiments were analyzed and plotted with Prism V9.0 (GraphPad, San Diego, CA, USA). Shown is the mean value + SEM unless indicated otherwise. Specific lysis was analyzed using two-way ANOVA with Sidak’s multiple comparison test. Cytokine and proliferation data were analyzed as indicated in the respective figure legends.

#### B cell culture

B cells were purified from PBMCs of healthy donors using negative isolation with immunomagnetic beads (Miltenyi Biotec) and subsequently expanded for 10-14 days in RPMI-1640 supplemented with 10% human serum (Bavarian Red Cross), 2mM Glutamax (Gibco, ThermoFisher), 100 U/ml penicillin–streptomycin (Gibco, ThermoFisher), using CD40L-multimere-kit and IL-4 (Miltenyi Biotec) according to manufacturer’s instructions.

#### Tumor cell culture

JeKo-1, Jurkat, Raji, MDA-MB-231 (all from the German collection of microorganisms and cell cultures (DSMZ), Braunschweig, Germany), and K562 (ATCC, Manassas, VA, USA) cells and the respective sublines were maintained in RPMI-1640 medium containing 8% fetal calf serum (FCS) (Sigma, F7524), 2 mM L-glutamine, and 100 U/mL penicillin/streptomycin (both Gibco, ThermoFisher) at 37°C and 5% CO_2_. Cell splitting was carried out according to the recommendation of the ATCC. ffluc/GFP-positive sublines and K562 or MDA-MB231 sublines expressing transgenes were generated by lentiviral transduction with the respective vectors with an MOI of 3 (see above).

#### Flow Cytometry

T cells were stained with one or more of the following conjugated mAbs: CD4 (clone SK3), CD8 (SK1), CD3 (Hit3a) and matched isotype controls (all BioLegend, San Diego, CA, USA). CAR-transduced (thus EGFRt+) T cells were detected by staining with anti-EGFR antibody (clone AY13, Biolegend, San Diego, CA, USA) and scFv-Linker (custom made, evitria, Schlieren, Switzerland). Staining with 7-AAD (BD, Franklin Lakes, NJ, USA) was performed for live/dead cell discrimination as directed by the manufacturer. Samples were measured on a FACS Canto II (BD Franklin Lakes, NJ, USA) or MACSQuant (Miltenyi Biotec, Bergisch Gladbach, Germany) and data analyzed using FlowJo software V.9.0 or V.10.0 (Treestar, Ashland, OR, USA).

#### Staining for *d*STORM analysis

For microscopy measurements Cellvis chamber slides (8 Chambered Coverglass Sytem #1.5 High Performance Cover Glass (0.17±0.005), Cellvis) were coated with 0.1 mg/mL poly-D-lysine (PDL) after which the cells were seeded into the chambers and centrifuged for adherence. Afterwards, the cells were kept on ice for 5 minutes, washed with HBSS, and stained with an antibody directly conjugated to Alexa Fluor 647 (AF647) at a concentration of 5 μg/mL. Subsequently, the cells were washed, fixed and imaged. The latter set of ROR2-specific CAR-T cells as well as the CARs with different linker lengths and spacer/transmembrane domains were stained in suspension and adhered on a CD45-antibody coating to reduce unspecific binding to the PDL.

#### *d*STORM imaging

For reversible photoswitching of AF647, a PBS-based imaging buffer (pH 7.4) was used that contained 100 mM β-mercaptoethylamine (Sigma-Aldrich). *d*STORM measurements were performed as previously described^3, 33–35^. Typically, 15,000 frames were recorded with a frame rate of ∼100 Hz (10 ms exposure time).

#### *d*STORM image reconstruction and data analysis

The recorded *d*STORM images were reconstructed with rapidSTORM 3.3 ^54^. Localization data acquired in *d*STORM measurements were analyzed using LOCAN, a custom-made code^3^. For analysis of each *d*STORM image an appropriate region of interest at the basal membrane of the cell, was chosen. For clustering analysis, a DBSCAN clustering algorithm was applied to group detected localizations^55^. Suitable parameters were ε = 20 and minPoints = 3. To distinguish between small and big localization clusters an arbitrary localization threshold of 40 was used, localization clusters with less than 40 localizations were counted as small, whereas localization clusters with more than 40 localizations were considered large localization clusters. In each experiment at least 15 cells were analyzed.

#### Preparation of co-cultures for microscopy

For the co-culture samples, K562_CD19^+^_CD20^+^ were mixed with ZAP70-GFP expressing CAR-T cells, ZAP70-GFP CAR-Jurkat cells, or Jurkat cells expressing a GFP-tagged CD19-CAR in a 1:10 ratio. As a control, ZAP70-GFP expressing cells were used that were not equipped with a CAR construct. The cells were then seeded into a PDL-coated Cellvis chamber and cultured for 30 minutes at 37°C and 5% CO_2_. Afterwards, the cells were stained as described for *d*STORM analysis with antibodies against scFv-Linker (evitria) or CD20 antibody (clone 2H7, Biolegend). Samples were stored at 4°C until imaging.

#### Airyscan imaging

Imaging was performed on a ZEISS LSM 900 with airyscan 2 using optimal pinhole and z-sectioning size. To achieve a good signal to noise ratio, each frame was scanned twice and signals were added. For the red channel a 640nm laser at 1% laser power, the orange a 561nm with 1.8% laser power and the green a 488nm laser were used. Due to the different expression levels of ZAP70-GFP and CAR-GFP, different settings for each condition were chosen. Namely, imaging was performed using 2% or 4% laser power and a pixel dwell time of 1.15µs or 2.30µs for ZAP70-GFP and CAR-GFP, respectively.

#### Quantification of accumulation at the contact site

CAR-Jurkat cells were labeled with Cellpose segmentation AI and an estimated diameter of 300 px. Clusters were also segmented with Cellpose^56^ using an estimated diameter of 15 px. Due to the different topology tumor cells were labeled with Stego^57^ segmentation AI. The area of interest is the boundary between tumor cells and Jurkat cells. To identify this area, Jurkat cell and tumor cell masks were converted to binary images. All entries in the Jurkat cell image, positioned under the tumor cell mask were set to zero. Binary dilatation was applied to the tumor cell mask image enlarging the mask size by one pixel. Subsequently summing the Jurkat cell masks and the enlarged tumor cell masks yields a value of two for pixels on the boundary. These boundaries are used for further processing. A mask for the cluster quantification at the boundary site was computed by applying a binary dilation of 25 px on the boundaries. Subsequently, clusters were counted, and this count was normalized on the covered area. The resulting value is compared with the clusters per area of Jurkat cells that are not in contact with a tumor cell. Each Jurkat cell - Jurkat cell to tumor boundary is treated as a separate value in the statistic. Values are plotted in a box plot. Significance is computed with Dunn’s significance test since the values are not normally distributed.

#### Combined SIM/*d*STORM imaging

Co-cultures were prepared as described above. Imaging was performed in *d*STORM imaging buffer on a ZEISS Elyra 7 with Lattice SIM². First, a three-color z-stack of 10 slices with 91 nm z-difference was taken of the contact site (red: 642nm, 1% laser power, 100 ms exposure time, orange: 561nm, 1% laser power, 80 ms exposure time, green: 488nm, 2% laser power, 80 ms exposure time) followed by *d*STORM imaging of the basal membrane. Channel alignment was achieved by measuring tetraspecs before the start of imaging. SIM images were processed using the SIM^2^ algorithm in ZEN. Afterwards, the microscope was switched to wide-field mode. Images were taken using the 642nm laser at 99% laser intensity in TIRF mode. The fluorescence emission was filtered through a LBF 405/488/561/642 and collected on the sCMOS camera. Images of 15000 frames were acquired with 20 ms exposure time. Image reconstruction and data analysis were performed the same as the other *d*STORM images.

#### Live-cell lattice light-sheet imaging

For imaging of immunological synapse formation with primary ZAP70-GFP-expressing T cells, Raji cells or donor-matched B cells were centrifuged at 100 × g for 5 minutes and resuspended in the antibody staining solution. The staining was done with a CD20-antibody conjugated to ATTO643 diluted in live-cell imaging medium (RPMI1640 + 10% human serum). Pulsing of Raji cells with 100 µg/mL Staphylococcal Enterotoxin E (SEE, abcam) was done at least 30 minutes before staining. The cells were washed once and seeded into Cellvis chamber slides that were coated with a BCMA antibody (Raji cells) or a CD20 antibody (primary B cells). T cells were centrifuged and resuspended in live-cell imaging medium for imaging and added to the well after which acquisition was started.

Lattice light-sheet imaging was performed using the commercially available microscope Lattice light-sheet 7 (Carl Zeiss AG, Jena). Excitation of samples were carried out using a water-immersion objective lens ((at ∼30° angle to sample holder) of 13.3× magnification and numerical aperture of 0.40. Dedicated laser lines of excitation wavelengths (λ_exc_) of 488 and 647 nm were used for illuminating primary ZAP70-GFP (CAR-)T cells and ATTO643-labeled Raji and B cells respectively. For our measurements, we used 3% and 5% of total laser power respectively for 647 and 488 nm excitation sources. We used a light-sheet of ∼1 µm thickness and 30 µm length for the experiments. For image acquisition, an exposure time of 20 ms was chosen. We acquired images of 2048 × 2048 pixels and collected 1001 slices at an interval of 0.200 µm. Emission light was collected through a separate objective lens orthogonal to the excitation objective lens (placed at 60° angle to cover glass) of 44.83× magnification and N.A. of 1.0. Fluorescence emission was passed through emission filter combination of a band-pass 495-550 and a long-pass 655 which was finally collected by a sCMOS camera (Hamamatsu ORCA-Fusion sCMOS camera, Shizuoka, Japan). We performed image analysis using the Zen 3.7 (Blue edition) software. Briefly, we processed the acquired raw (skewed) LLS images as following: the images were deconvolved using the constraint iterative algorithm with 12 iterations. Deconvolution was followed by deskewing of the images, linear interpolation, and rotation to actual cover-glass coordinates. Supplementary movies provided were created with Imaris and Fiji (ImageJ) Microscopy Image Analysis Software (Oxford Instruments).

#### Reproducibility

All experiments were performed at least three times. Representative images are shown for each experiment.

#### Reporting Summary

Further information on research design is available in the Nature Research Reporting Summary linked to this article.

---

**Extended Data Fig. 1.**
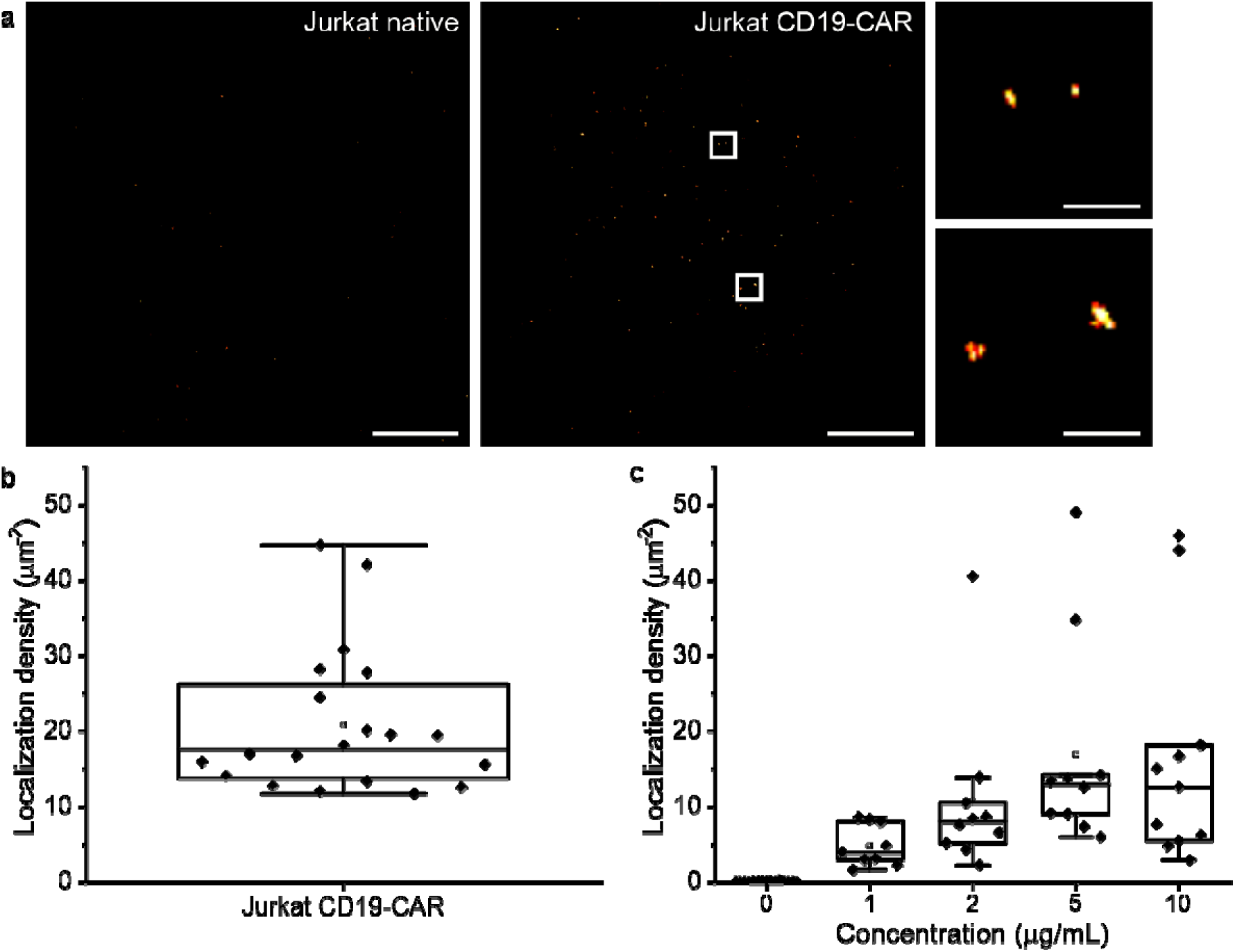
Visualizing expression and distribution of CD19-specific CARs in the plasma membrane of Jurkat cells. **a**, *d*STORM images show CAR localization clusters of varying size on the basal membrane of a CD19-specific CAR expressing Jurkat cell (right) but no expression on a native Jurkat cell (left). Small panels display magnification of boxed regions indicating that CARs are organized as monomers (top) and in CAR clusters (bottom). **b**, Boxplot of localization densities measured on CD19-specific CAR-T cells (n=17) with a median density of 17.62 ± 4.5 localizations µm^-2^. **c**, Localization densities detected on CD19-specific Jurkat cells at different antibody concentrations (n=8-10) show saturation at an antibody concentration of 5 µg ml^-1^. Scale bars, 3 μm (a), 300 nm (magnified images).

**Extended Data Fig. 2.**
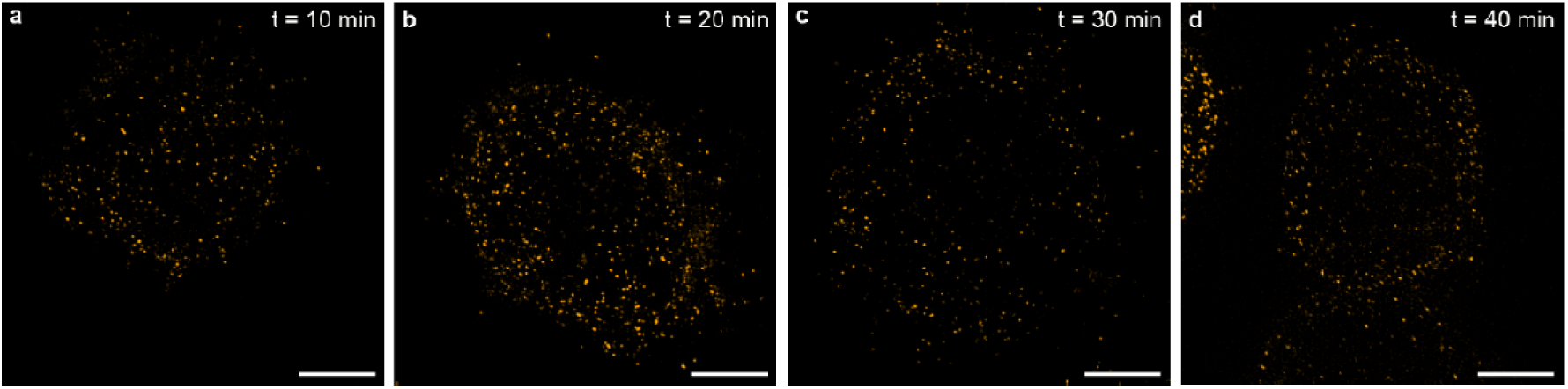
Antibody staining does not induce CAR nanodomain formation. **a-d**, SIM images (maximum z-projections) of CD19-specific CAR expressing Jurkat cells stained with primary antibody with different incubation times (10-40 min) before fixation and imaging. CAR nanodomains are visible already after 10 min antibody incubation time and do not change at longer incubation times suggesting that CAR nanodomains are not formed by antibody crosslinking. Scale bars, 5 μm (a-d).

**Extended Data Fig. 3.**
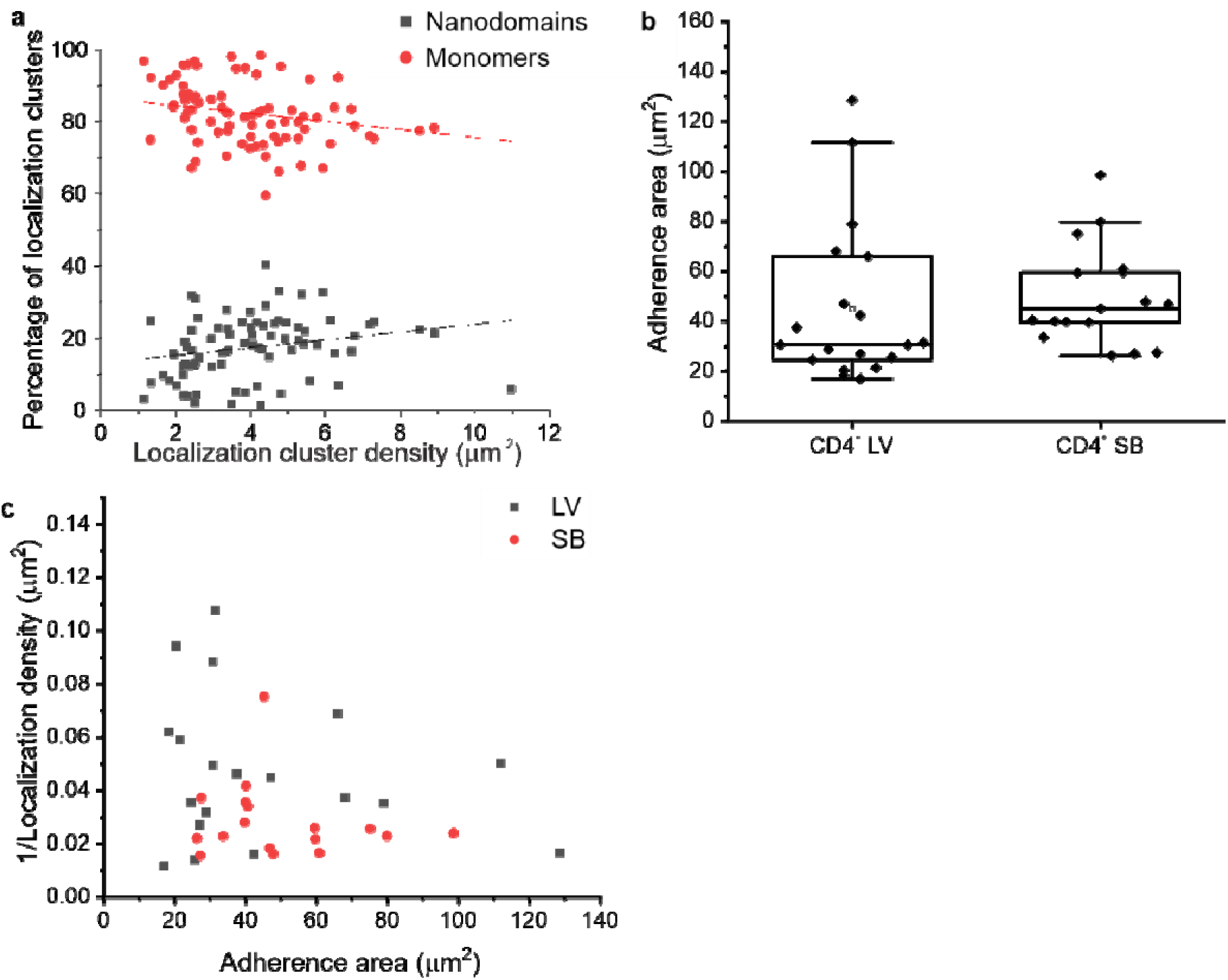
**a**, Percentage of CD19-specific CAR localization clusters detected as monomers (gray) and nanodomains (red) on CD4^+^ T cells with increasing overall CAR expression (n=82 cells). **b**, Boxplots show adherence areas (µm^2^) of CD4^+^ ROR2-specific CAR-T cells generated by LV transduction and SB MC transfection. **c**, Scatter plot of the inverse of the localization density (µm^2^) versus the adherence area (µm^2^) showing no relationship between localization density and adherence area.

**Extended Data Fig. 4.**
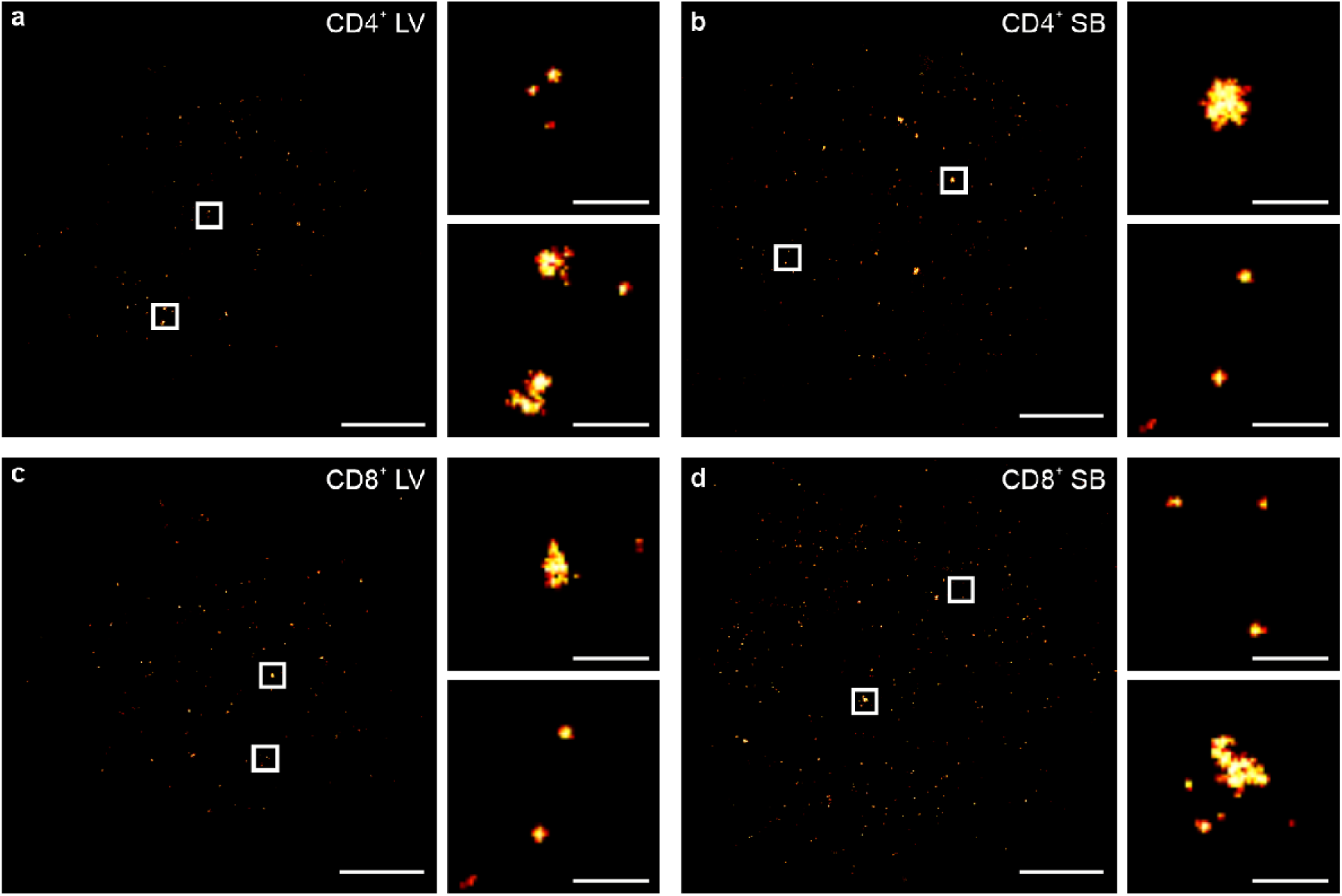
ROR2-specific CARs form nanodomains in the plasma membrane of T cells. **a,b**, *d*STORM images of ROR2-specific CARs on the basal membrane of primary CD4^+^ T cells introduced by lentiviral transduction and SB transfection. **c,d**, *d*STORM images of ROR2-specific CARs on the basal membrane of primary CD8^+^ T cells introduced by lentiviral transduction and SB transfection. Small panels display magnifications of boxed regions revealing the existence of CAR nanodomains in the plasma membrane independent of the used introduction method. Scale bars, 3 µm (a-f), 300 nm (magnified images).

**Extended Data Fig. 5.**
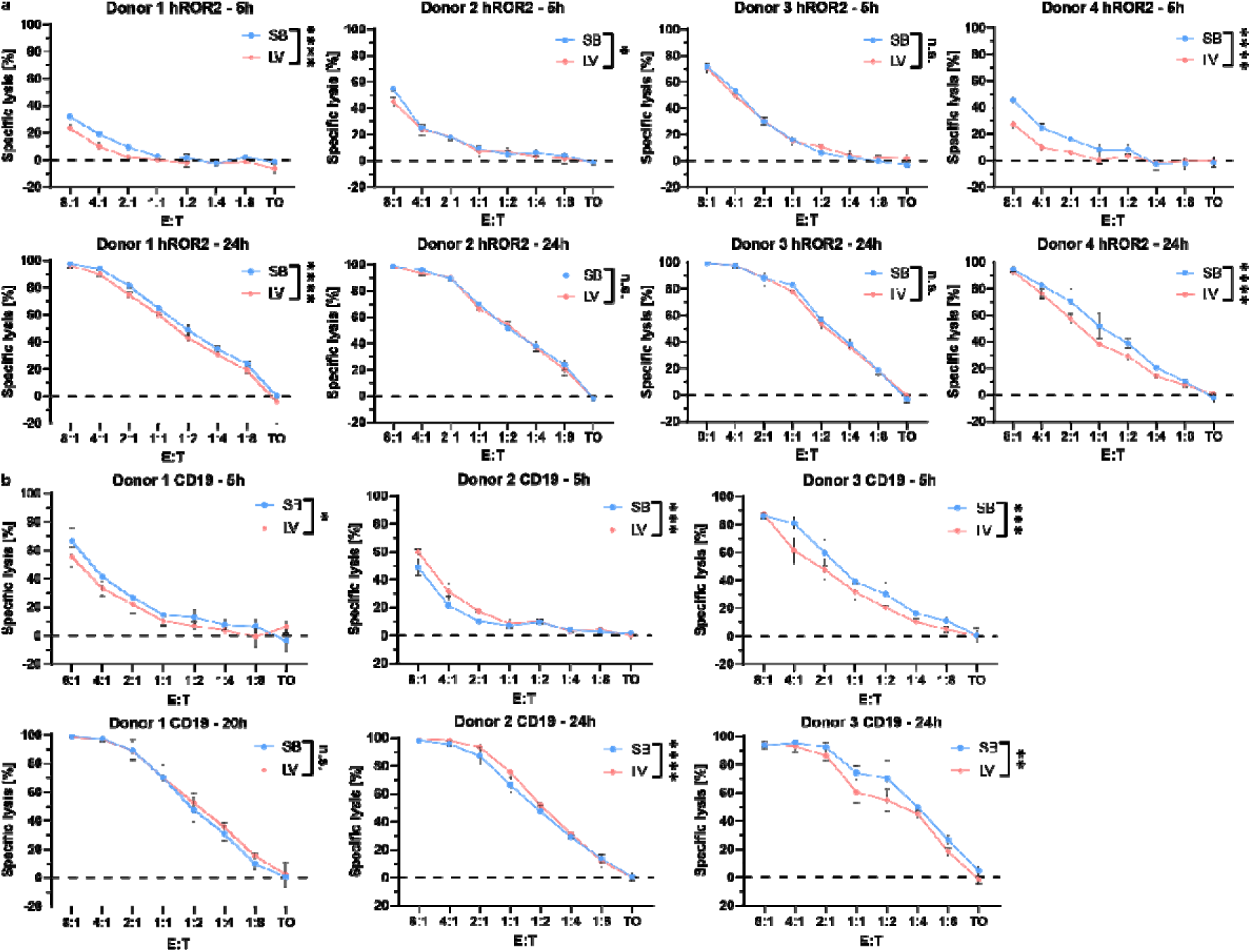
Short and long-term cytotoxicity of SB and LV CAR-T cells against MDA-MB-231 hROR2 and K562 CD19. Functional *in vitro* characterization of CAR-T cells generated from separate donors by Sleeping Beauty transposition (SB) or lentiviral transduction (LV), targeting either hROR2 or hCD19. Short (5h, upper panels) or long-term (24h, lower panels) antigen-specific lysis mediated by CD8^+^ CAR-T cells co-cultured with **a**, MDA-MB-231 (hROR2^+^) or **b**, or K562 (CD19^+^) in in multiple effector-to-target ratios (E:T) in luminescence-based cytotoxicity assays. Data are depicted as Mean ± SD of 3 technical replicates. Statistical analysis was performed using two-way ANOVA with Sidak correction for multiple comparisons. *p< 0.05, **p< 0.01, ***p< 0.001, ****p< 0.0001.

**Extended Data Fig. 6.**
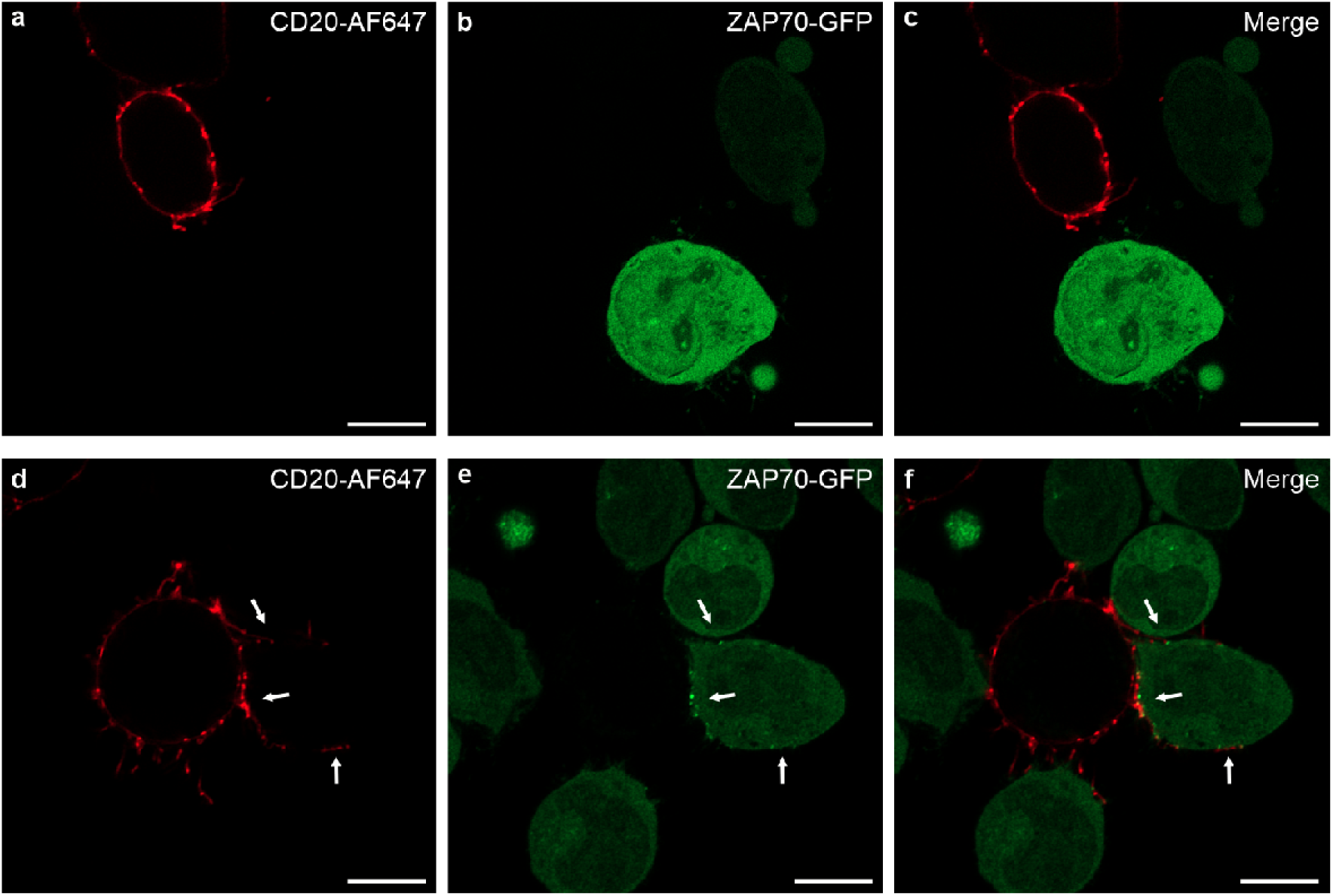
ZAP70 accumulates at contact sites of CD19-specific CAR-Jurkat cells and tumor cells. **a-c**, Confocal airyscan images of K562_CD19^+^_CD20^+^ and Jurkat cells expressing ZAP70-GFP co-cultured at a ratio of 1:10 for 30 min before immunolabeling with anti-CD20 antibodies and fixation. **d-f**, Confocal airyscan images of K562_CD19^+^_CD20^+^ and CD19-specific CAR-Jurkat cells expressing ZAP70-GFP co-cultured at a ratio of 1:10 for 30 min before immunolabeling with anti-CD20 antibodies and fixation. White arrows indicate contact sites between the cells highlighting ZAP70-GFP accumulation. Scale bars, 5 µm.

**Extended Data Fig. 7.**
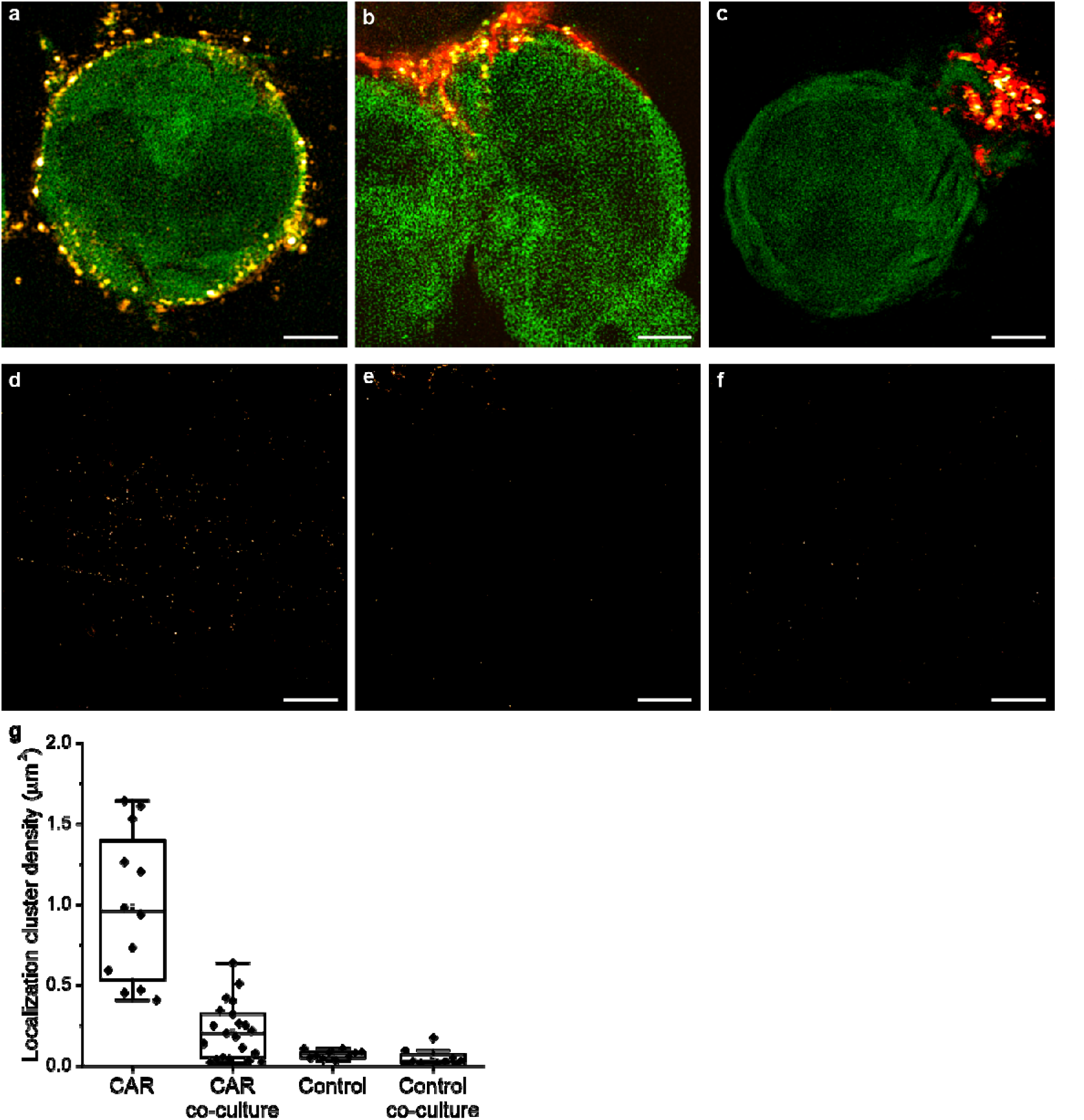
*d*STORM images of CD19-CARs in the absence and presence of tumor contacts. **a-d,**. Lattice SIM images of the CD19-specific CAR-Jurkat cells in the absence (a) and presence (b,c) of K562_CD19^+^_CD20^+^ tumor cells. Orange CAR-AF647, red, CD20-CF568 green ZAP70-GFP. **d-f**, Corresponding *d*STORM images of CD19-specific CAR-Jurkat cells depicted in a-c. **i**, Boxplots of localization cluster densities in the absence and presence (co-culture) of tumor cells indicating a significant reduction in CD19-CAR localization cluster density (monomers and nanodomains) upon contact formation with tumor cells. These data support the finding that CAR monomers and nanodomains move to the contact site. Scale bars, 3 µm.

**Supplementary Fig. 1.**
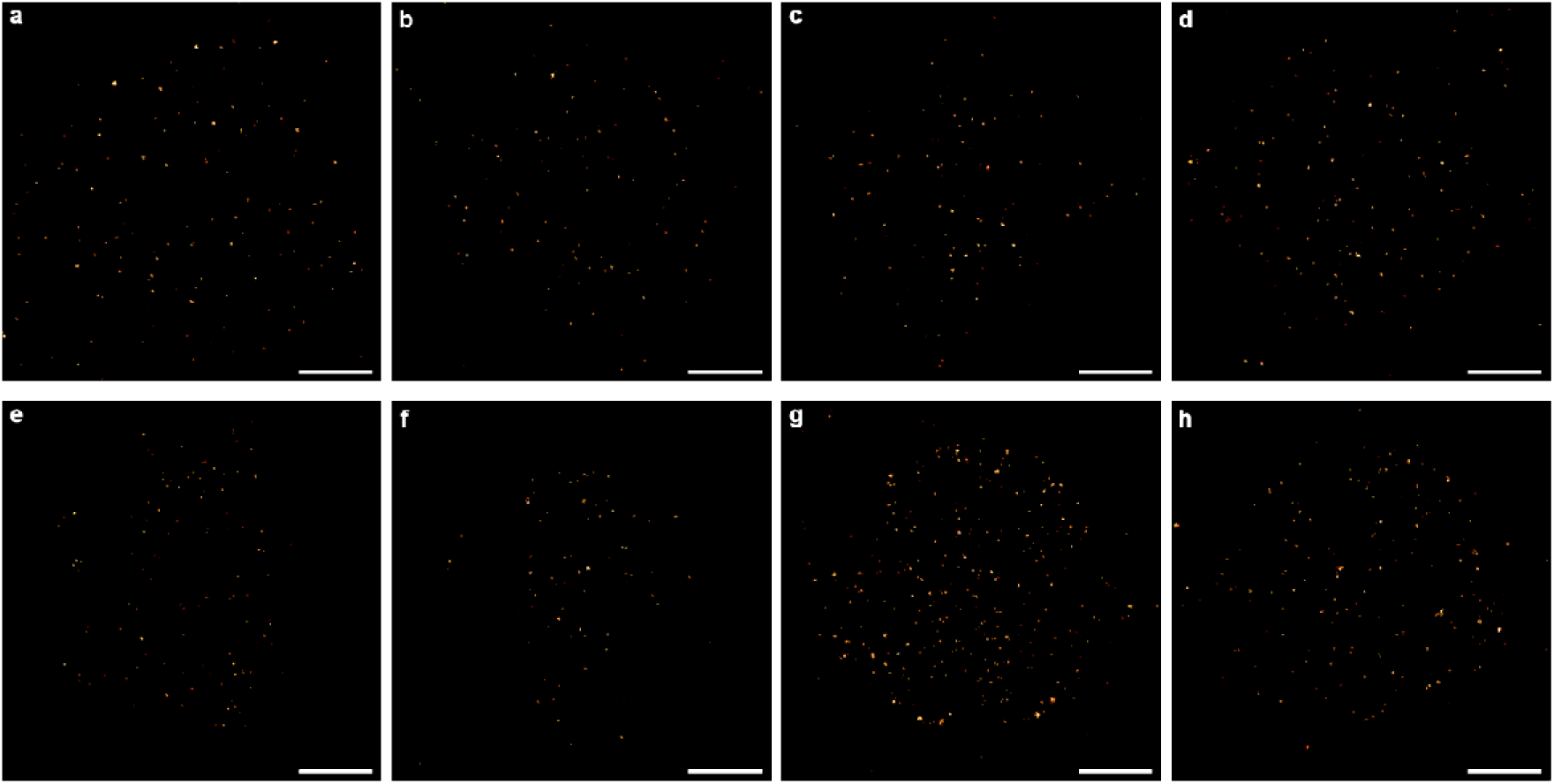
**a-d**, Example *d*STORM images of CARs on the basal membrane of CD19-specific CAR-Jurkat cells. **e-h**, *d*STORM images of CARs on the basal membrane of CD19-specific CAR-Jurkat cells recorded at different AF647-labeled antibody concentrations of 1 µg ml^-1^ (e), 2 µg ml^-1^ (f), 5 µg ml^-1^ (g), 10 µg ml^-1^ (h). Scale bars, 3 µm.

**Supplementary Fig. 2.**
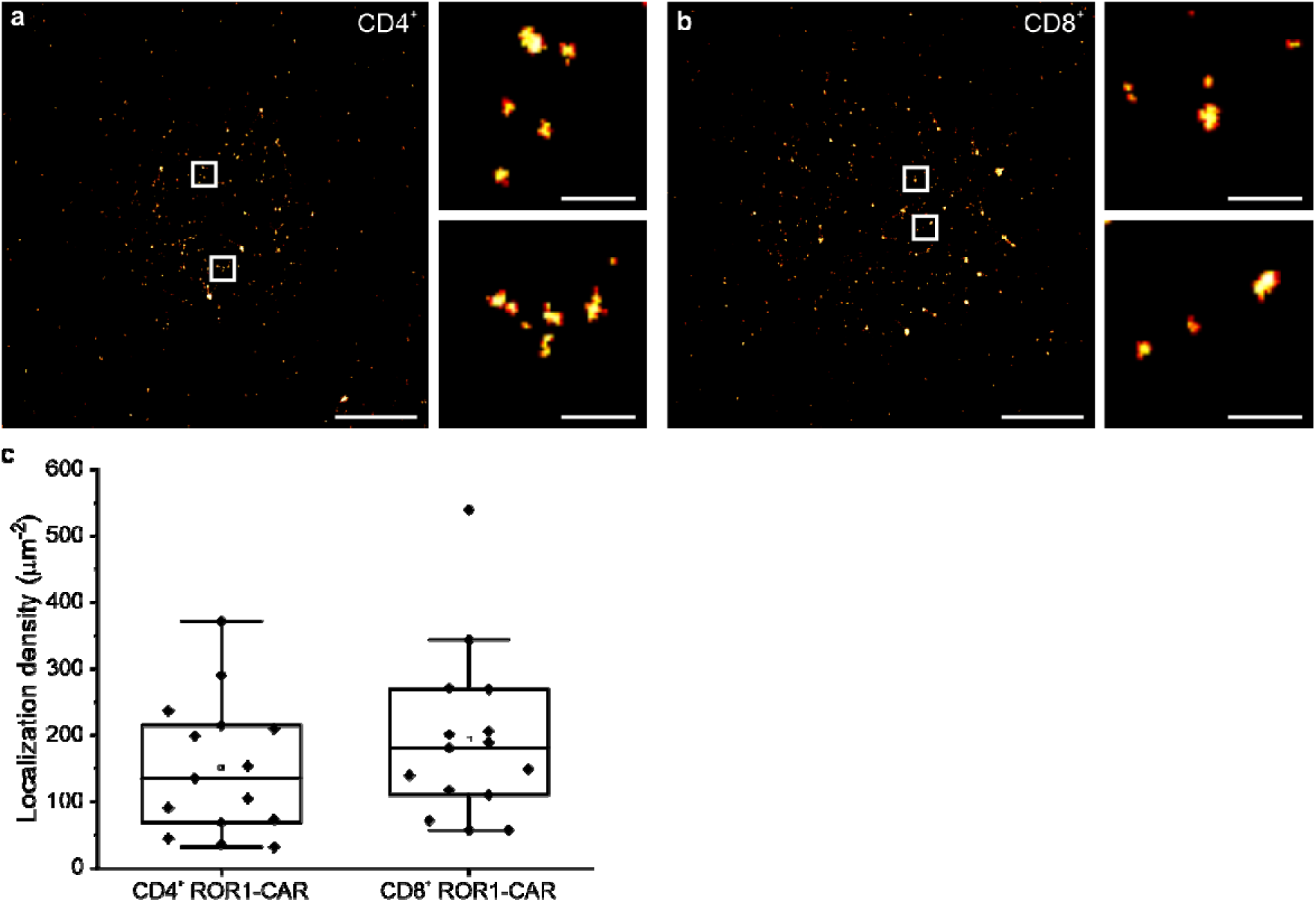
**a, b**, *d*STORM images of the basal membrane of CD4^+^ and CD8^+^ T cells equipped with ROR1-specific CARs. Small panels display magnifications of boxed regions revealing the organization of CARs as monomers and in nanodomains in the plasma membrane independent of the T cell phenotype. **c**, Boxplots show localization densities (localizations µm^-2^) of 135 ± 74 (CD4^+^), and 181 ± 71 (CD8^+^) for ROR1-specific CARs (n=15 cells). Scale bars, 3 μm (a, b), 300 nm (magnified images).

**Supplementary Fig. 3.**
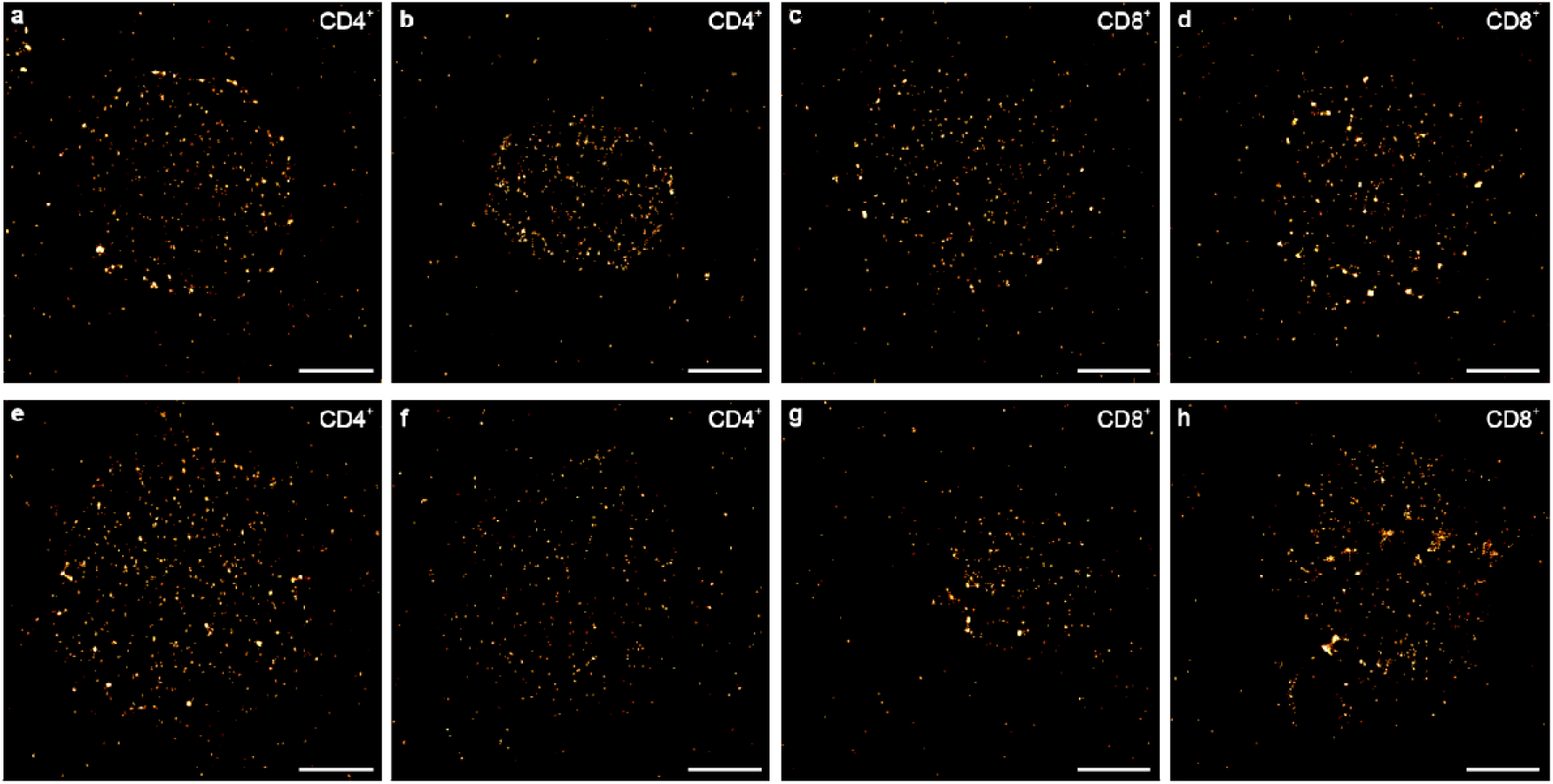
Example *d*STORM images of CD19-(a-d) and ROR1-specific CARs (e-h) on the basal membrane of primary CD4^+^ and CD8^+^ cells introduced by SB transfection. Scale bars, 3 µm.

**Supplementary Fig. 4.**
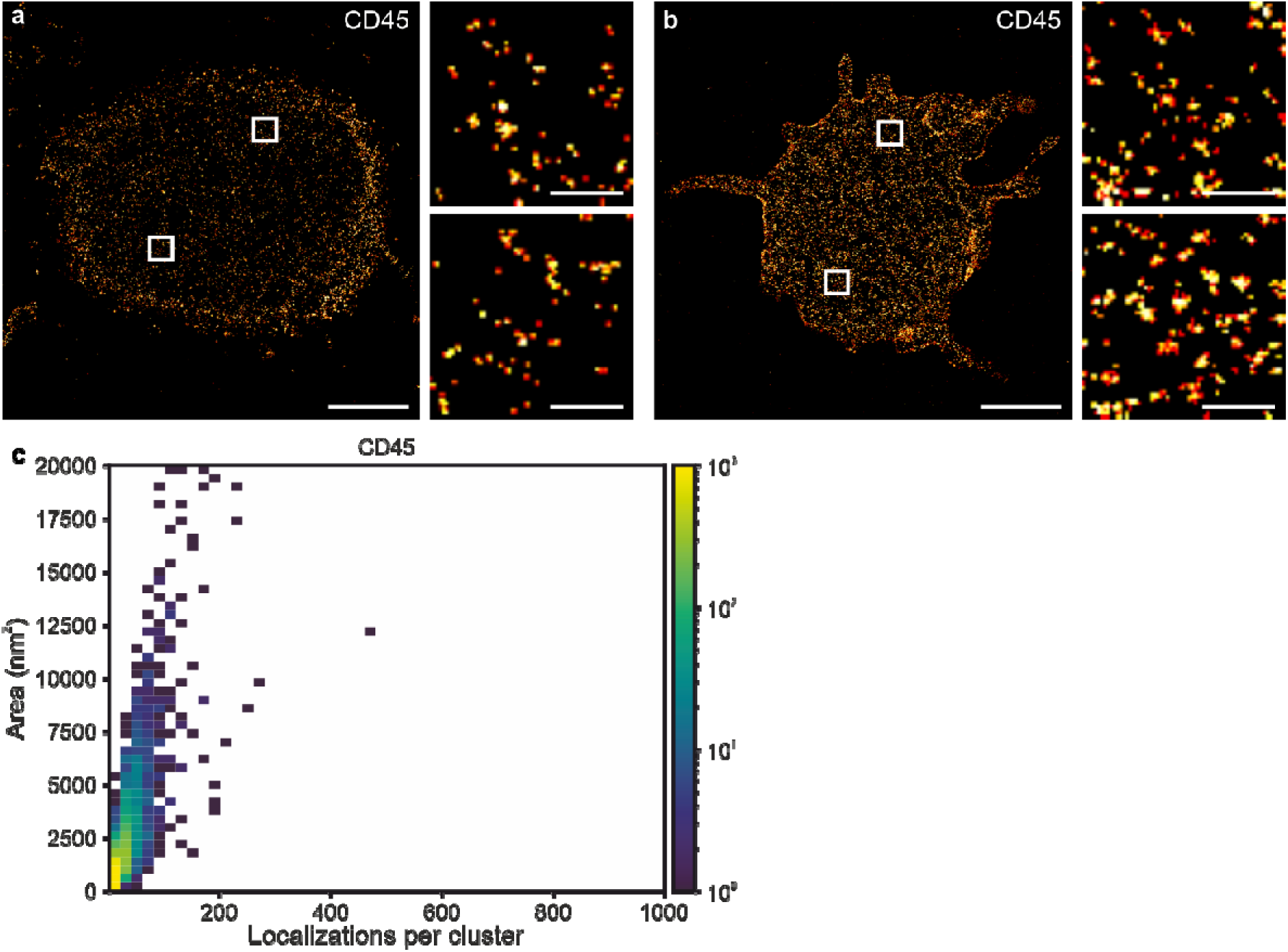
**a, b** *d*STORM images of CD45 in the plasma membrane of Jurkat cells (a) and CD4^+^ primary T cells (b) labeled with primary AF647 labeled antibodies. Scale bars, 3 µm, 300 nm (magnified images). **c** Bivariate histogram from spatially unresolvable localization clusters in *d*STORM experiments of CD45 in the plasma membrane of Jurkat cells using AF647-labeled antibodies. Histograms display the distribution of localizations per cluster and convex hull areas of localization clusters.

**Supplementary Fig. 5.**
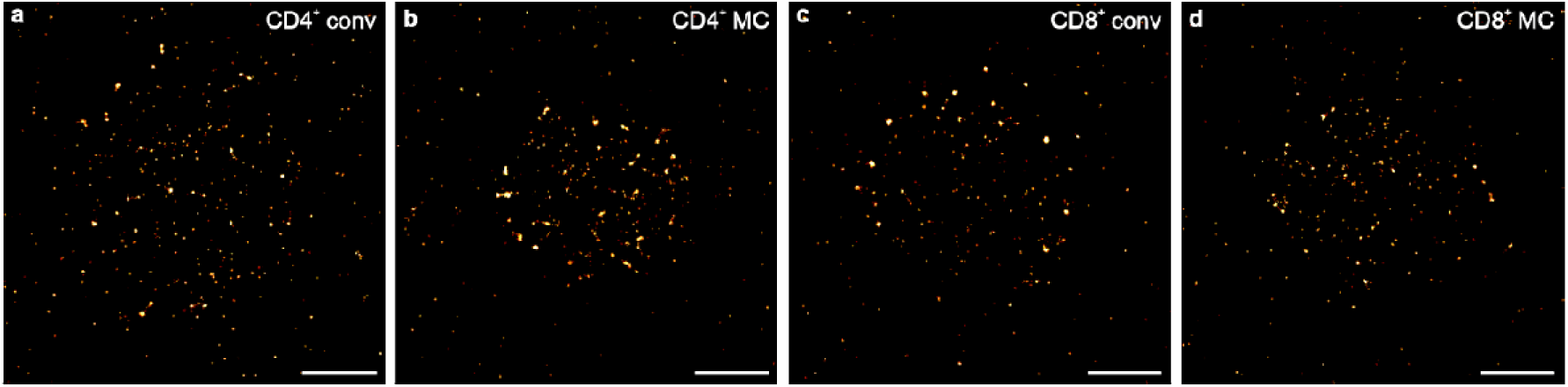
**a-d**, Example *d*STORM images of CD19-specific CARs on the basal membrane of primary CD4^+^ (a, b) and CD8^+^ (c, d) cells introduced by conventional SB plasmids (conv) and SB minicircle (MC). Scale bars, 3 µm.

**Supplementary Fig. 6.**
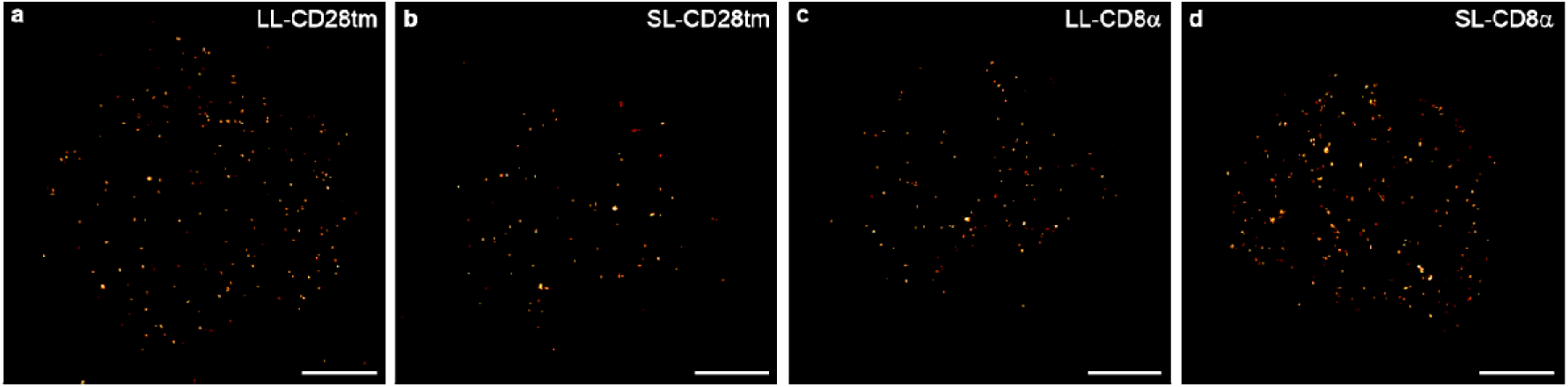
**a-d**, Representative *d*STORM images of CD19-specific CAR-T cells with different transmembrane domains and linker lengths. **a,b**, IgG spacer and CD28-based transmembrane domain **b,c**, a CD8α-based spacer and transmembrane domain. The scFv-linker was shortened for CAR constructs shown in images **b** and **d**. Scale bars 3 µm.

**Supplementary Fig. 7.**
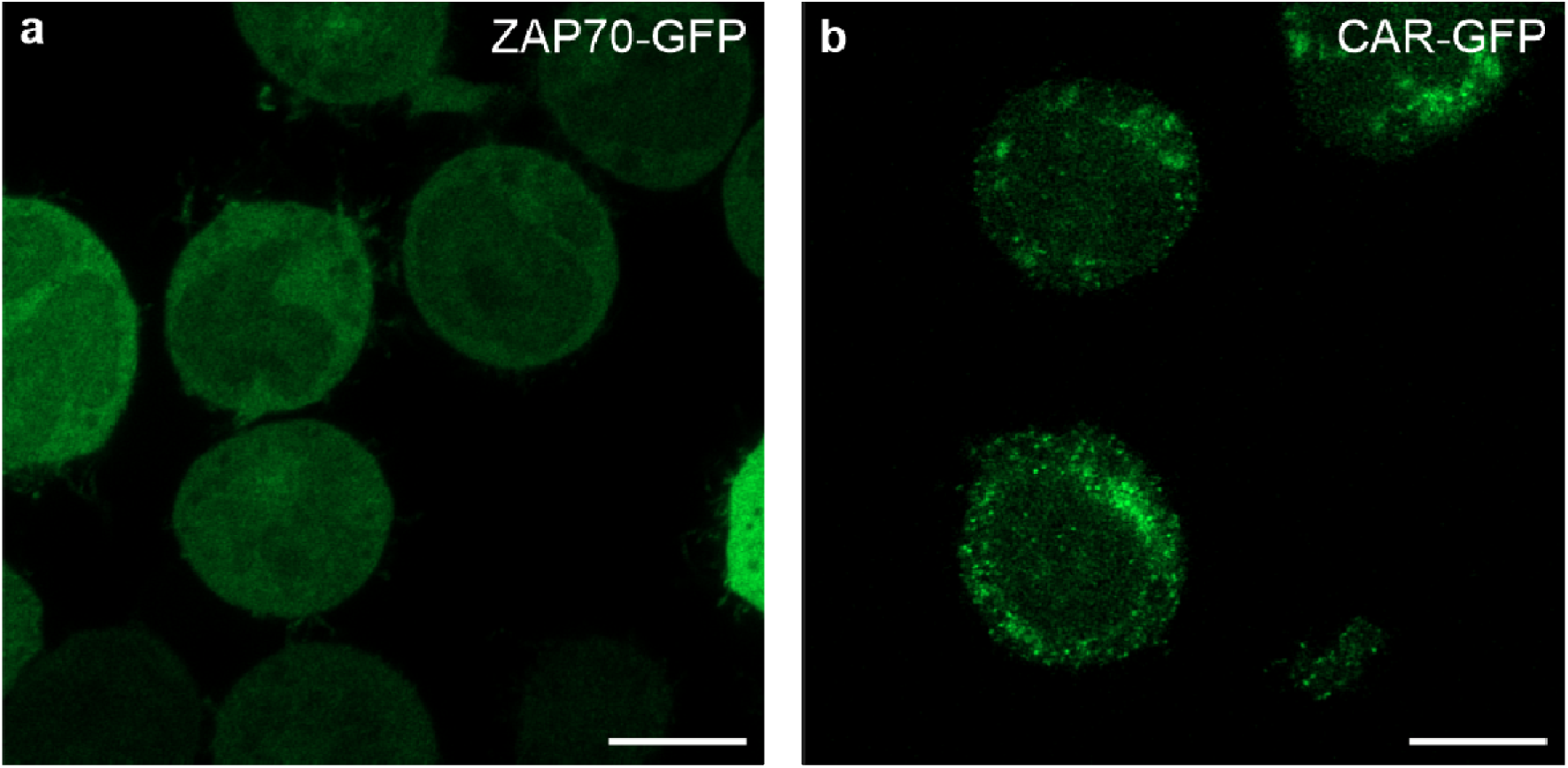
**a,** Confocal airyscan fluorescence image of CD19-specific CAR-Jurkat cells expressing ZAP70-GFP displaying a homogenous distribution of ZAP70 in the absence of tumor cells. **b**, Confocal airyscan fluorescence image of CAR-Jurkat cells expressing CD19-specific GFP-tagged CARs. The image shows that CARs are distributed over the entire plasma membrane in the absence of tumor cells. Scale bars, 5 µm.

**Supplementary Fig. 8.**
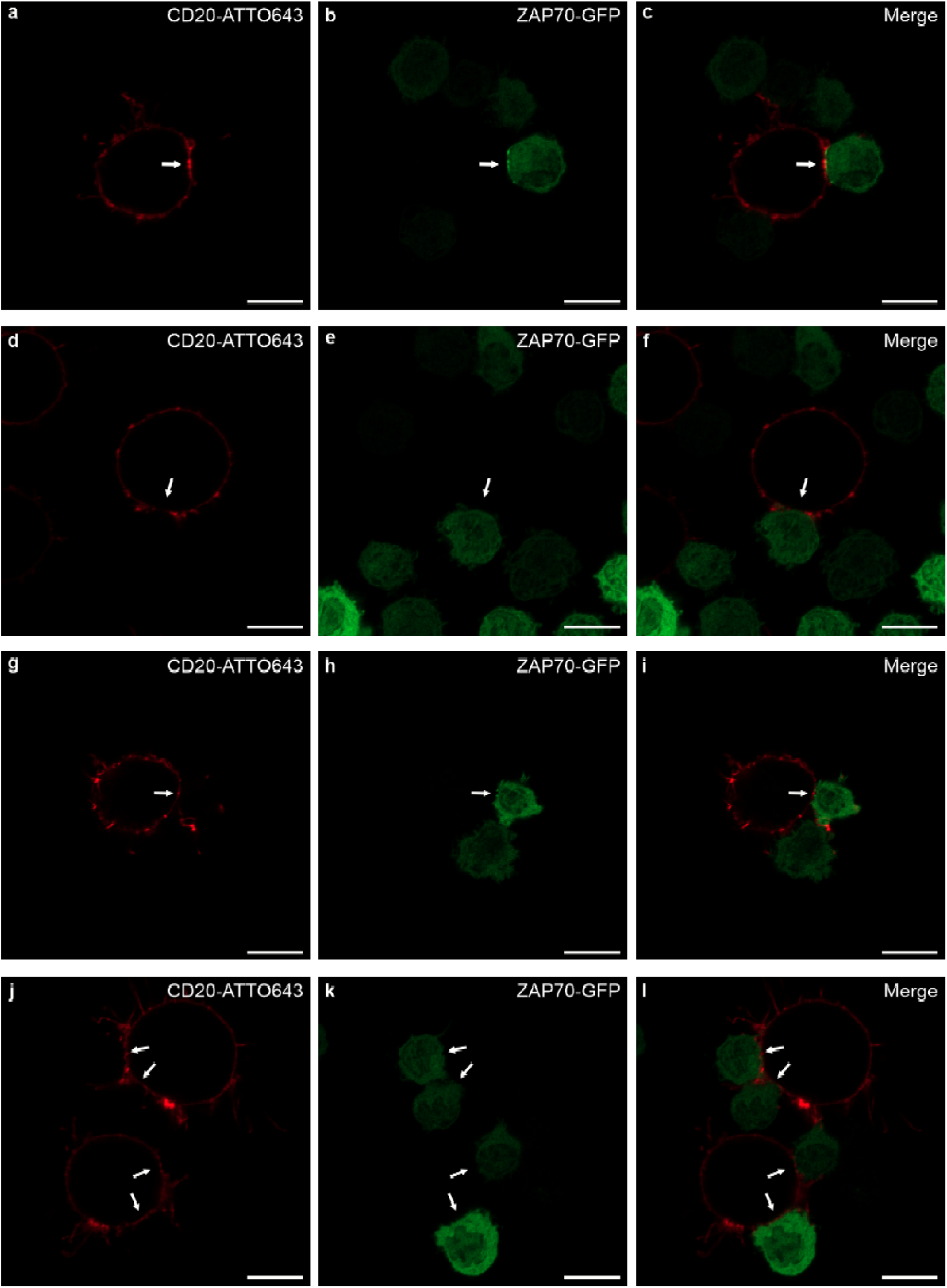
ZAP70 accumulates at contact sites of CD19-specific CAR-T cells and tumor cells. **a-f**, Confocal airyscan images of K562_CD19^+^_CD20^+^ and CD4^+^ T cells expressing ZAP70-GFP and CD19-CAR (a-c) or no CAR (d-f) co-cultured at a ratio of 1:10 for 30 min before immunolabeling with anti-CD20 antibodies and fixation. **g-l**, Confocal airyscan images of K562_CD19^+^_CD20^+^ and CD8^+^ T cells expressing ZAP70-GFP and CD19-CAR (g-i) or no CAR (j-l) co-cultured at a ratio of 1:10 for 30 min before immunolabeling with anti-CD20 antibodies and fixation. White arrows indicate contact sites. Scale bars, 5 µm.

**Supplementary Fig. 9.**
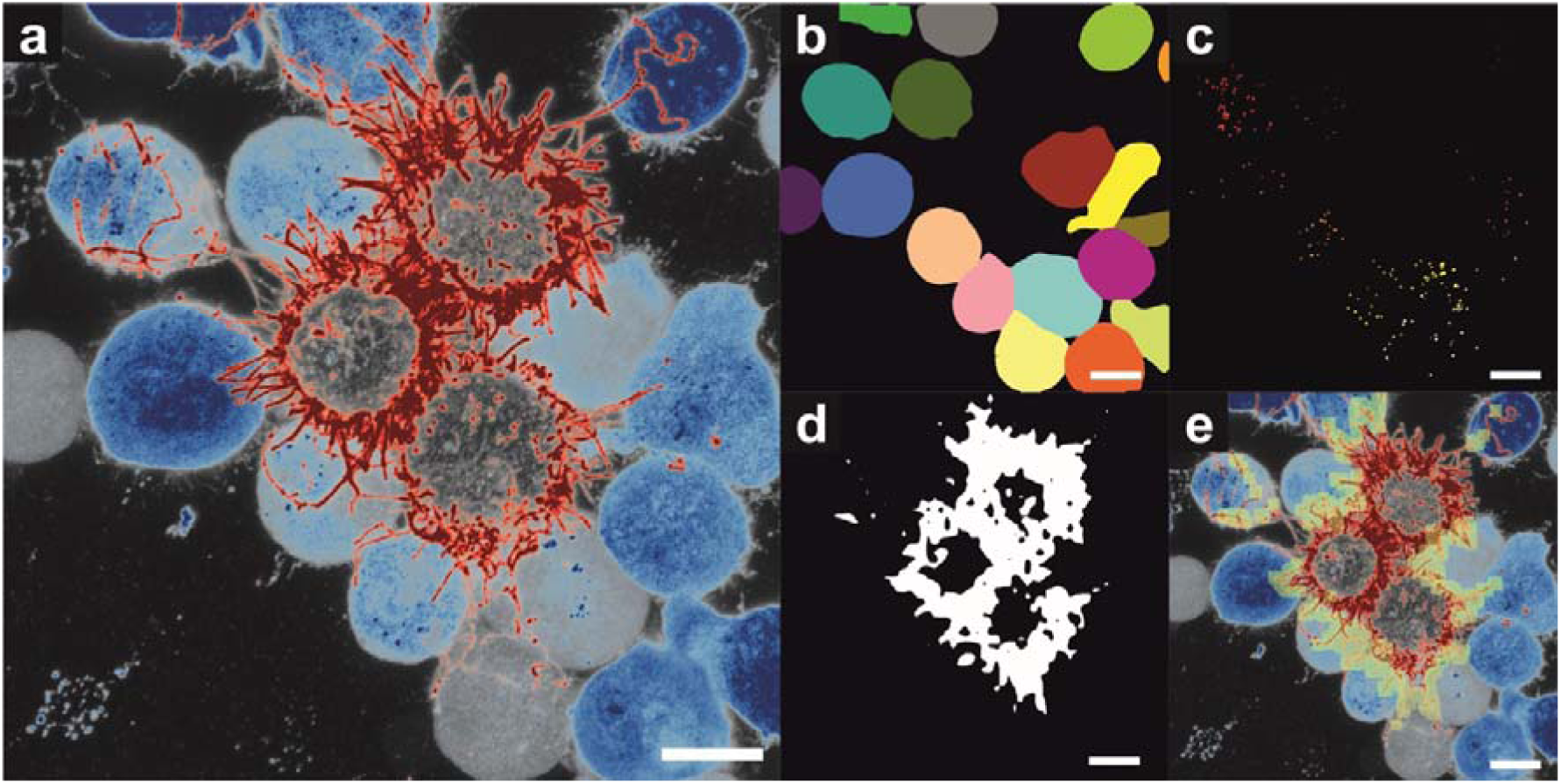
Data evaluation. **a**, CD19-specific CAR-Jurkat cells (blue) and tumor cells (red). **b**, CAR-Jurkat cells are labeled with Cellpose segmentation AI and an estimated diameter of 300 px. **c**, Intensity clusters are segmented with Cellpose using an estimated diameter of 15 px. **d**, Tumor cells are labeled with Stego segmentation AI. The area of interest is the boundary between tumor and Jurkat cells. We convert the Jurkat and the tumor cell masks to binary images and set all entries in the Jurkat cell image, positioned under the tumor cell mask to zero. We apply a binary dilatation to the tumor cell masks enlarging their size by one pixel. Subsequently summing the Jurkat cell masks and the enlarged tumor cell masks yields a value of two for pixels on the boundary. These boundaries are used for further processing. **e**, using an area around the boundary that covers 25 px in each direction (computed by dilation of the boundary line) we compute a mask for the cluster quantification at the boundary site. We subsequently count the clusters and normalize this count on the covered area. The resulting value is compared with the clusters per area of Jurkat cells that are not in contact with a tumor cell. Each cell/cell boundary is treated as a separate value in the statistic. Scale bars, 10 µm.

**Supplementary Fig. 10.**
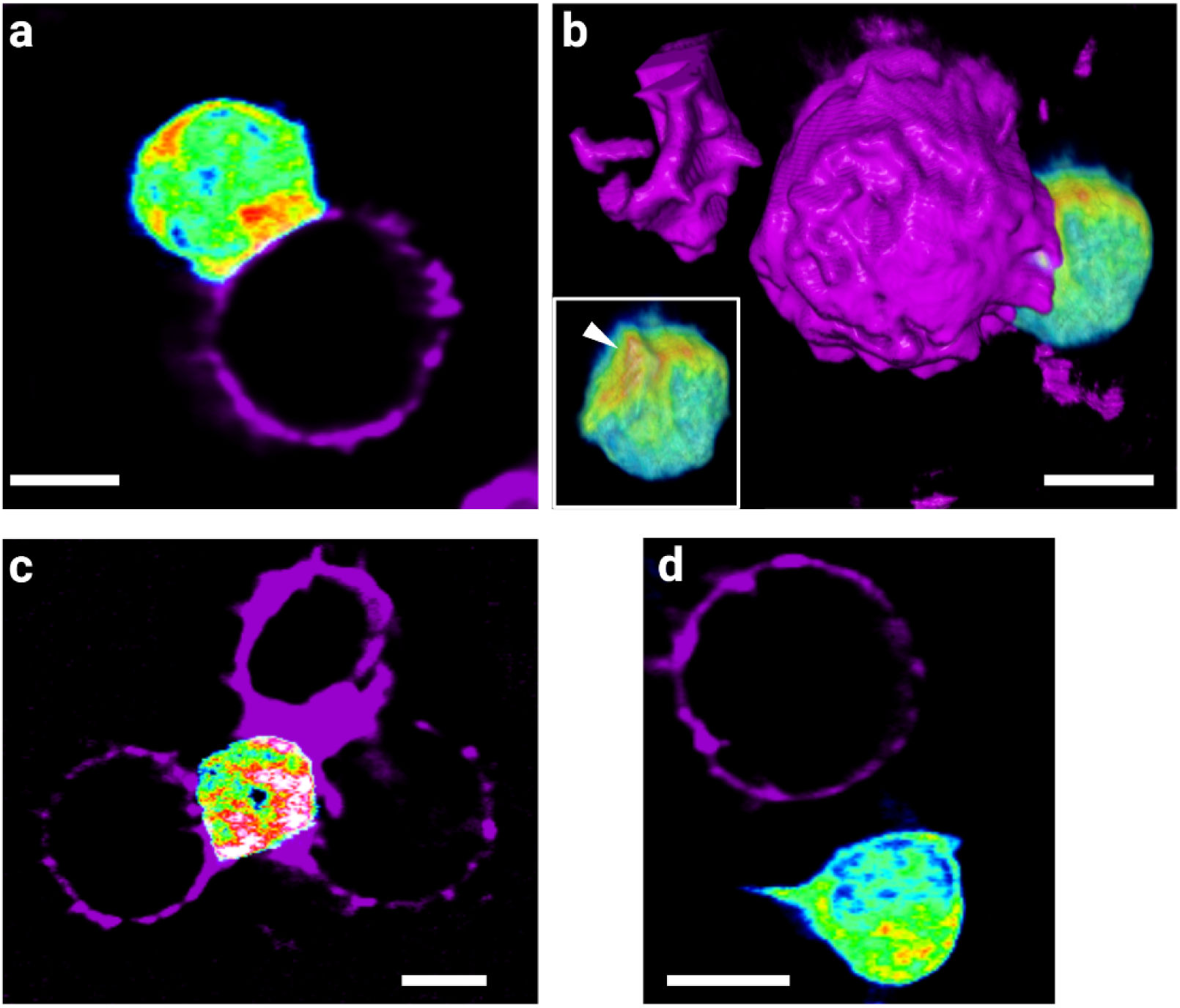
Lattice light-sheet imaging of immunological synapse formation with CD8^+^ T cells. **a,** Top view of a ZAP70-GFP expressing (rainbow color scale) CD19-specific CAR-T cell in contact with a Raji cell (magenta) stained with CD20-ATTO643 antibodies. **b,** Side view of the same contact site showing accumulation of ZAP70 signal at the contact site (indicated by white arrows). The insert shows that ZAP70 accumulates at the contact site. **c,** CD8^+^ ZAP70-GFP expressing T cell not equipped with a CAR forming a classical immunological synapse with SEE-pulsed Raji cells. **d,** Only rapid or no contacts are formed between CD8^+^ ZAP70-GFP expressing T cells when no CAR is present. Scale bars, 5 μm

**Supplementary Fig. 11.**
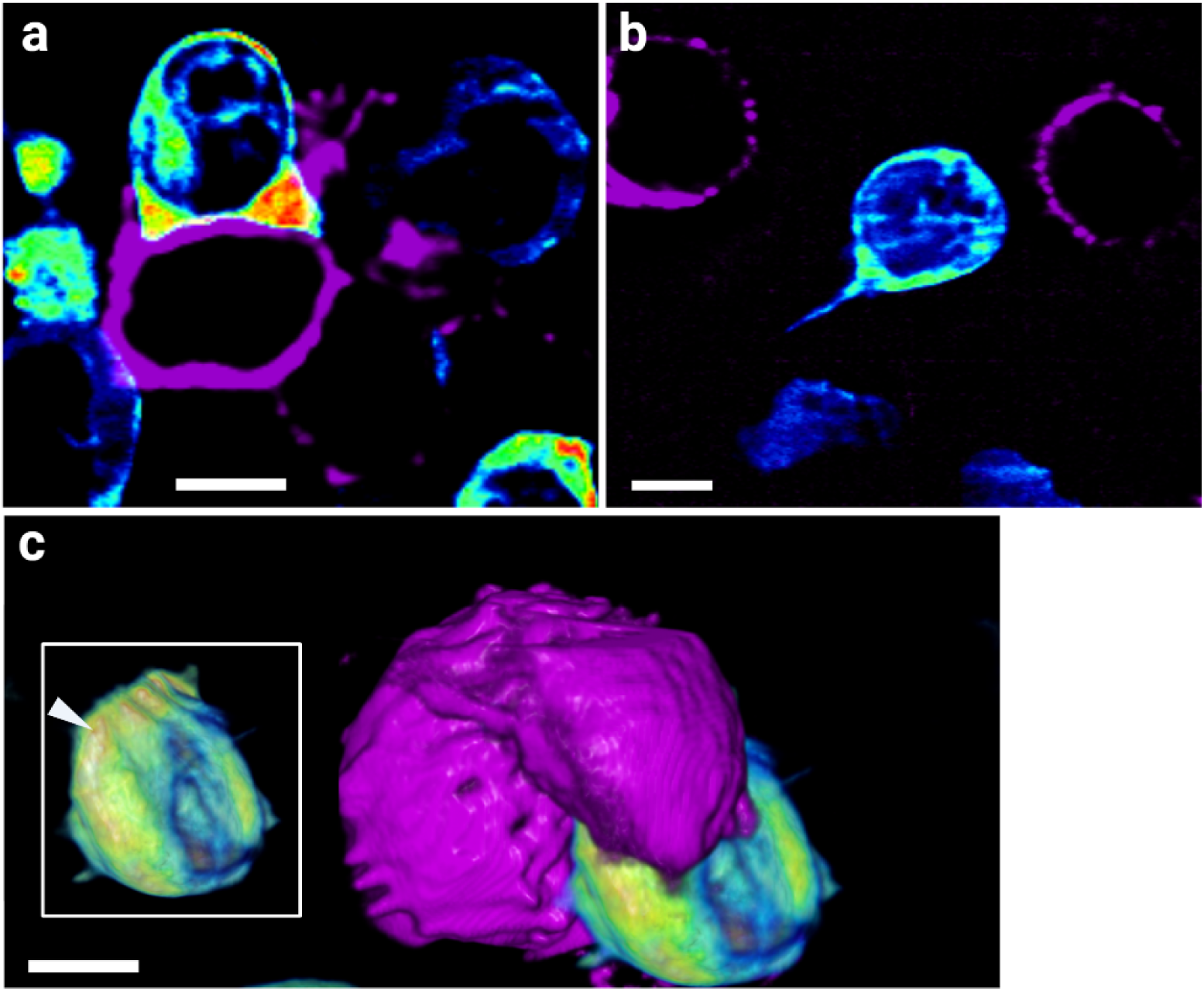
Lattice light-sheet imaging of immunological synapse formation between ZAP70-GFP expressing CD4^+^ T cells and donor-matched B cells. **a,** Top view of a ZAP70-GFP expressing (rainbow color scale) CD19-specific CAR-T cell in contact with a B cell (magenta) stained with CD20-ATTO643 antibodies. **b**, ZAP70-GFP expressing CD4^+^ T cells that were not equipped with a CD19-specific CARs formed no or very short contacts with B cells, without any indication of ZAP70 accumulation. **c,** Side view of the same contact site from (a) showing accumulation of ZAP70 signal at the contact site (insert shows the ZAP70-GFP channel only). Scale bars, 5 μm.

**Supplementary Fig. 12.**
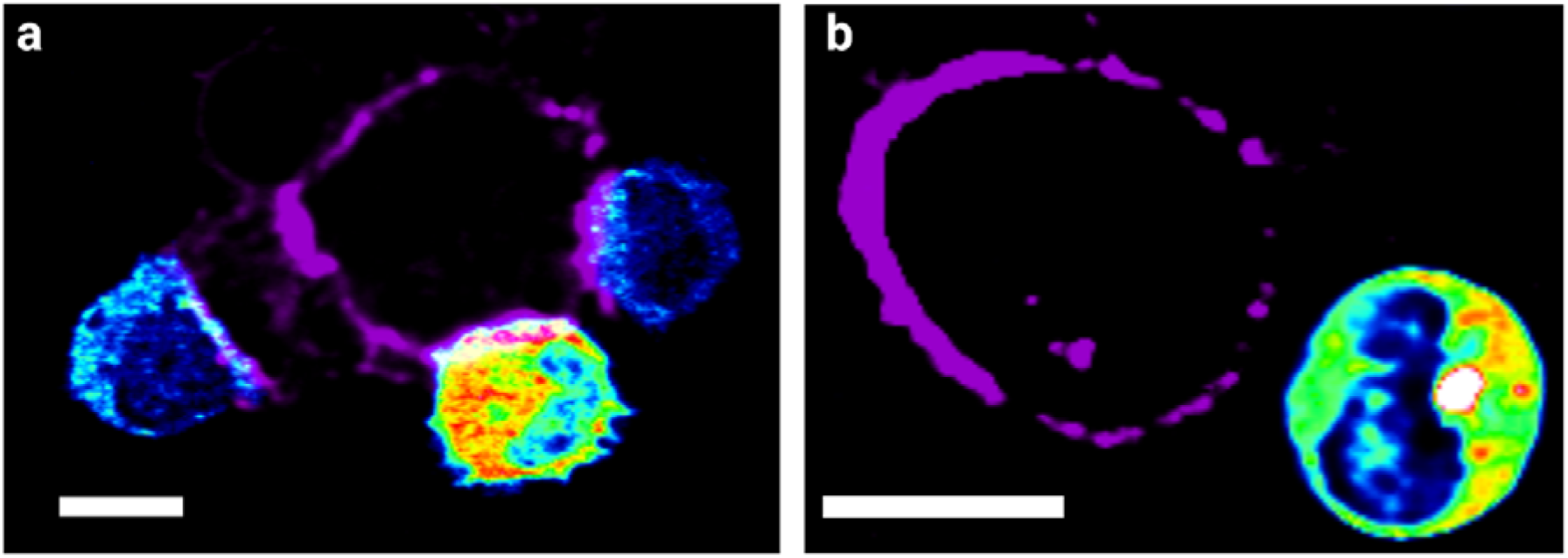
Lattice light-sheet imaging of immunological synapse formation between ZAP70-GFP expressing CD8^+^ T cells and donor-matched B cells. **a,** Top view of a ZAP70-GFP expressing (rainbow color scale) CD19-specific CAR-T cell in contact with a B cell (magenta) stained with CD20-ATTO643 antibodies. **b**, ZAP70-GFP expressing CD4^+^ T cells that were not equipped with a CD19-specific CAR formed no or very short contacts with B cells, without any indication of ZAP70 accumulation. Scale bars, 5 μm.

## Supplementary Movies

**Supplementary movies S1-S4:** Movies show exemplary *z* slices illustrating the formation of an immunological synapse (IS) between Raji cells immunostained with CD20-antibody ATTO643 (magenta) and CD4^+^ (S1, S2) or CD8^+^ CD19-specific CAR-T cells expressing ZAP70-GFP (rainbow LUT) (S3, S4). Relative times are annotated. Total acquisition time was 40 min. Movies are played at 5 fps.

**Supplementary movies S5, S6:** Movies show exemplary *z* slices illustrating the interaction between Raji cells immunostained with CD20-antibody AF647 (magenta) and CD4^+^ (S5) or CD8^+^ (S6) ZAP70-GFP T cells that do not express CD19-CARs (rainbow LUT). No stable contact formation followed by accumulation of GFP signal is seen, confirming CAR-induced clustering and stable IS formation. Relative times are annotated. Total acquisition time was 40 min. Movies are played at 5 fps.

**Supplementary movies S7-S8:** Movies show exemplary *z* slices illustrating the formation of an immunological synapse (IS) between donor-matched primary B cells immunostained with CD20-antibody ATTO643 (magenta) and CD4^+^ (S7) or CD8^+^ CD19-specific CAR-T cells expressing ZAP70-GFP (rainbow LUT) (S8). Relative times are annotated. Total acquisition time was 40 min. Movies are played at 5 fps.

**Supplementary movies S9-S10:** Movies show exemplary *z* slices illustrating the formation of a “bull’s eye”-shaped immunological synapse (IS) between Raji cells immunostained with CD20-antibody AF647 (magenta) and CD4^+^ (S9) or CD8^+^ ZAP70-GFP T cells (rainbow LUT) (S10) where Raji cells were pulsed with staphylococcal enterotoxin type E (SEE). Relative times are annotated. Total acquisition time was 40 min. Movies are played at 5 fps.

